# A conservation planning assessment of basin wide Unionid mussel assemblages using environmental DNA

**DOI:** 10.64898/2026.02.13.705757

**Authors:** Nathaniel Marshall, Megan Seymour, Nathan Herbert, Cheryl Dean, W. Cody Fleece

## Abstract

Conservation planning for rare, threatened, and endangered species requires basic information for distribution and abundance. Often this information is lacking due to the nature of traditional survey methods which can be time and labor intensive and thus costly. Environmental DNA (eDNA) metabarcoding offers a promising approach for monitoring freshwater mussel assemblages, a taxonomic group that is both highly imperiled and difficult to survey using traditional methods. We evaluated the performance of eDNA metabarcoding across 30 km of Fish Creek, in Ohio and Indiana, U.S.. We compared results to visual surveys conducted at the same sites. eDNA detected 25 mussel species, including four species not observed alive visually, while visual surveys detected 22 live species. Both methods confirmed the presence of three federally protected species, and eDNA uniquely detected *Simpsonaias ambigua*, a species rarely encountered in conventional surveys. Incorporating detection repeatability improved congruence between methods: high-repeatability detections strongly aligned with visual presence, whereas moderate and low repeatability detections likely represented reach-scale occupancy. Overall, eDNA metabarcoding offers an efficient and sensitive tool for assessing mussel assemblages and can substantially enhance monitoring programs when integrated with species ecology and hydrological context.

## Introduction

Freshwater mussels (order Unionida) represent one of the most imperiled faunal groups in North America. Within the United States, 303 mussel species are currently recognized (FMCS 2025), of which 75 are federally endangered and an additional 21 are federally threatened (USFWS 2025). Their rarity, combined with their tendency to burrow into benthic substrates, makes traditional survey methods labor-intensive and highly dependent on expert taxonomists (Smith 2006, Wisniewski et al. 2013, Sanchez & Schwalb 2019). Further, the lack of comprehensive data for distribution and abundance constrains efforts to assess status, historical trends, and overall recovery planning for rare, threatened, and endangered species. These challenges highlight the need for innovative approaches that can improve detection efficiency and accuracy to support effective conservation and management.

Environmental DNA (eDNA), genetic material shed into the environment through waste, mucus, gametes, or sloughed cells, has become increasingly integrated into natural resource monitoring programs aimed at detecting rare species and characterizing biological communities (Beng & Corlett 2020, Deiner et al. 2021). Environmental DNA sampling offers several advantages over conventional survey methods, including higher sensitivity, reduced cost, minimal habitat disturbance, and improved access to remote or difficult-to-sample habitats (Jerde 2021, Sternhagen et al. 2024). In particular, eDNA metabarcoding, which enables simultaneous identification of multiple species from a single environmental sample, is emerging as a promising tool for assessing mussel assemblages in lotic systems (Marshall et al. 2022, Marshall & Fleece 2025, Prié et al. 2025).

Several studies have demonstrated that eDNA approaches can be particularly effective for characterizing diverse mussel assemblages (Prié et al. 2021, Coghlan et al. 2021, Marshall et al. 2022). Because unionid communities often include species that vary widely in abundance, detectability, and habitat use, traditional surveys may underrepresent rare, cryptic, or deeply burrowed taxa (Smith 2006, Wisniewski et al. 2013, Sanchez & Schwalb 2019). Whereas eDNA may be able to capture a more complete representation of assemblage composition, enabling researchers to quantify diversity and community assemblages across large spatial scales (Dokai et al. 2023, Prié et al. 2025, Johnson et al. 2025). Environmental DNA-based assessments are poised to become a central tool for evaluating mussel biodiversity at both local and landscape scales (e.g., Marshall et al. 2022, Prié et al. 2025, Coghlan et al. 2021).

More than 70% of freshwater mussel species in North America are considered endangered, threatened, or of conservation concern (Haag & Williams 2014, Williams et al. 2017). Population declines have been driven by multiple stressors, including degraded water quality, habitat loss, declines in host fish populations, and the spread of invasive species (Williams et al. 1993, Ricciardi et al. 1998). Mussels play critical ecological roles in freshwater ecosystems by filtering water, cycling nutrients, modifying benthic habitats, and serving as prey for other organisms (Gutiérrez et al. 2003, Vaughn 2018).

Fish Creek in northwestern Ohio and northeastern Indiana, US, supports a diverse assemblage of native freshwater mussels (Watters 1998). Over the past few decades, Fish Creek has seen a noticeable decline in mussel richness (Watters 1998, 2000). Of particular interest, Fish Creek is the last known extant location of *Epioblasma perobliqua* (White Cat’s Paw Pearly Mussel), which has not been observed since 1999 (Watters 2000), even though surveys have been conducted within the system since (USFWS 2020, 2023).

This study implemented eDNA metabarcoding and visual surveys to assess the distribution and abundance of rare and endangered freshwater mussels. This project used eDNA and visual surveys to (1) provide a description of stream wide mussel assemblage within Fish Creek, (2) assess site-level eDNA detection of known mussel assemblages and evaluate the level of effort required to detect rare species, and (3) compare spatial estimates of species occupancy from eDNA and visual data. Both surveys were implemented to help assess the current status of *E. perobliqua* and other species of conservation concern within Fish Creek.

## Methods

### Visual Mussel Survey

Visual surveys were conducted using low-effort qualitative methods to maximize the spatial extent of surveyed sites across Fish Creek. The objective was to rapidly survey as many sites as possible rather than to exhaustively survey any single site. These qualitative surveys were intended to determine if formerly identified mussel beds persist, and document spatial diversity and mussel density patterns. At each site, biologists examined the substrate and identified if suitable habitat for mussels was present. Once suitable habitat was determined, biologists flagged the survey starting point. Each search started at the downstream end of the site and moved in an upstream direction.

Searchers formed a line across the width of the stream and moved steadily upstream while searching, essentially forming parallel transects that covered the majority of the stream bed over the search area. Surface searches were performed, which included moving cobble and woody debris; hand sweeping away silt, sand, and/or small detritus; and disturbing/probing at least the upper 5 cm of loose substrate to better observe mussels. Viewing buckets were used at several sites where depth and water clarity permitted.

Search effort was haphazard between sites for a number of reasons. First, the number of searchers varied according to participant availability. Second, each search lasted for 20 minutes regardless of the number of searchers. Third, the total length and area of the search varied by site according to site conditions, number of mussels encountered, and number of searchers. Therefore, at each site, the search area (m^2^) and search effort (total people hours) were recorded. The search area widely varied across sites, ranging from 38 m^2^ to 330 m^2^ (mean 134 ± 72 m^2^). The search time varied across sites, ranging from 1.67 to 6.67 search hours (mean 3.5 ± 1.4 hours) (Supplementary Table 1).

Surveys were conducted within the late summer over three sampling years (2021-2023). Surveys were completed on September 9 through September 10 in 2021, August 31 through September 1 in 2022, and August 21 through August 22 in 2023 (Table 1). The visual surveys consisted of 13 sites in 2021, 14 sites in 2022, and 12 sites in 2023 (Table 1, Figure 1). Survey sites spanned from near the confluence of the Saint Joseph River (0.16 kilometers) to 28 kilometers upstream of the confluence (Figure 1).

**Figure 1.**
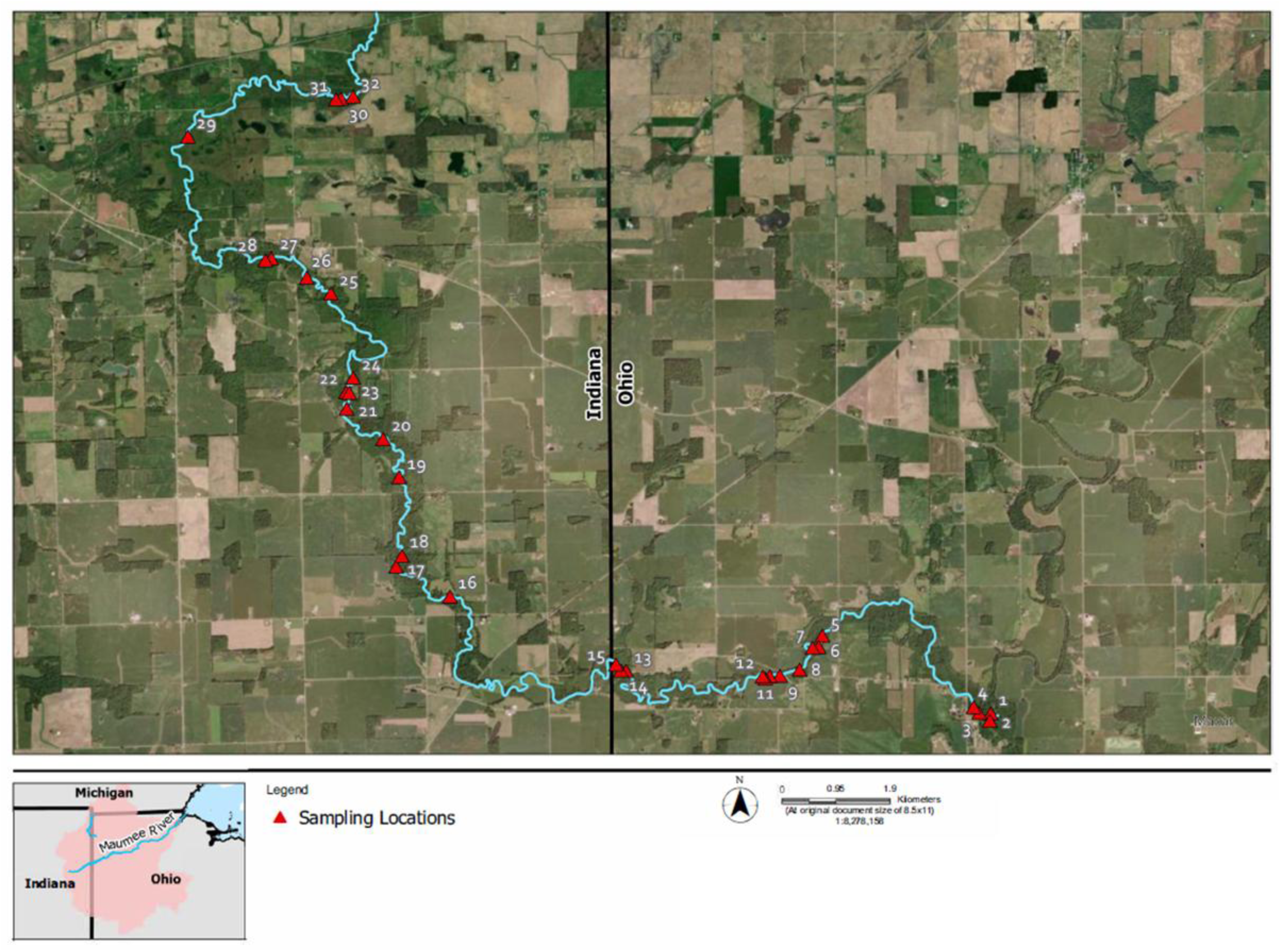
Map of eDNA survey sites sampled in Fish Creek.

**Table 1.**
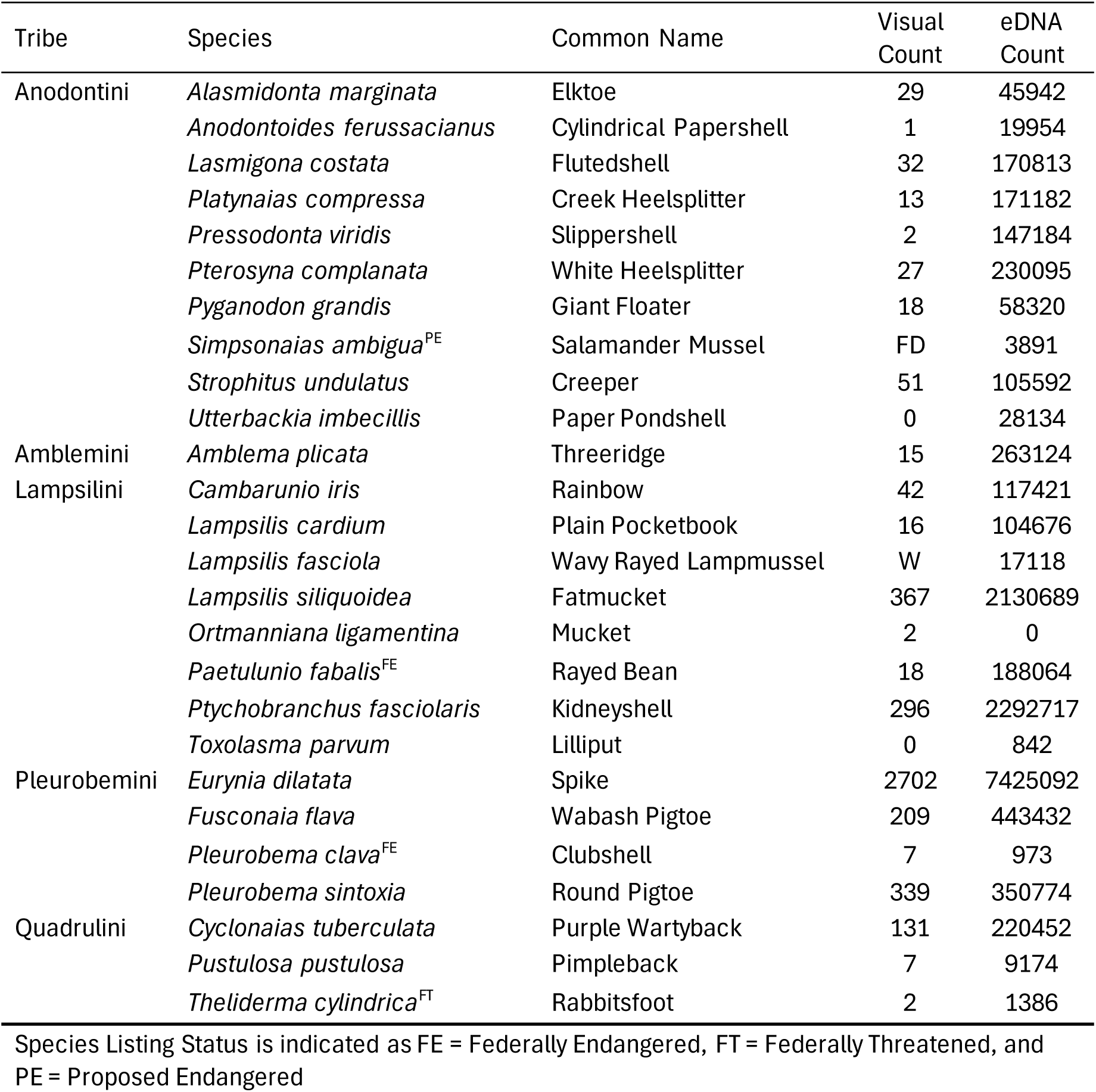
Species surveyed during the visual and eDNA sampling events in Fish Creek. FD = fresh dead shell, W = weathered shell.

### Environmental DNA Survey

#### Sample Collection

The eDNA sampling was always conducted prior to any visual surveys to avoid sediment disturbance during sampling. A total of 121 eDNA water samples were collected during the 2021, 2022, and 2023 Fish Creek sampling events (Table 1). An additional 15 negative field and/or filtration controls were collected across the three sampling events (five in 2021, five in 2022, and five in 2023). The eDNA survey included all sites searched during the visual survey (Table 1), with an additional site near the confluence of the Saint Joseph River and three additional sites upstream near river mile 18 (Table 1). Due to the close proximity of some sites within the visual survey, mussel data from some visual survey sites are grouped together as a single eDNA site (Table 1). The subsequent search area (ft ^2^) and search effort (people hours) were likewise summed for visual survey sites that were grouped together (Table 1).

In 2021, the eDNA was collected as replicate point samples across the width of the creek. Three 500mL point samples were collected so that the first was collected near one bank, the second near the middle of the creek, and the third from the opposite bank. Each 500mL sample was collected in a clean Nalgene container and then placed in a dark cooler on ice. Upon leaving the field, the water was filtered from each sample using a 47-mm-diameter glass microfiber filter GF/C (nominal pore size 1.2µm; GE Healthcare Life Science, MA, USA). After vacuum-pumping, filters were placed into separate coin envelopes, which were then placed in Ziploc bags with silicone desiccant beads and stored in a freezer (−20°C). On each of the two field collection days, negative field controls were collected on average every 2 to 3 sites. This was accomplished by opening a Nalgene bottle of 500mL of distilled water in the field, and then processing it alongside the field samples. Additionally, a filtration negative control was collected each day, by filtering 500mL distilled water alongside field samples.

In 2022 and 2023, 500-1000mL water samples were collected from near the benthos using a peristaltic pump with a polypropylene filter holder attached to a painter’s pole. eDNA was collected along a transect spanning the width of the river for each replicate sample. Once the first sample was collected, the surveyor took the next sample ∼2-3m upstream, and continued this for at least three replicates per site. Samples were filtered on a 47-mm-diameter GF/C. Filters were preserved the same as in 2021. On each field collection day, negative field controls were collected on average every 2 to 3 sites. This was accomplished by filtering 500mL of distilled water from a clean Nalgene bottle in the field using the same peristaltic pump system.

All collection equipment was decontaminated with 10% bleach solution after each day of sampling. The eDNA surveyor wore new sterile nitrile gloves at each site and when handling eDNA equipment.

### Laboratory Processing

All filters were shipped on ice to Cramer Fish Science Genidaqs (Sacramento, California, USA, https://genidaqs.com) for DNA extraction and metabarcoding using Illumina MiSeq processing.

#### Sample DNA Extraction

The DNA was extracted from each filter using a modified Qiagen DNeasy® Blood and Tissue Kit protocol. Each filter was processed overnight at 56 °C in 540 µl ATL and 60 µl Proteinase K. The resulting supernatant was passed through a Qiashredder spin column, mixed with 600 µl AL and incubated at 70 °C for 10 min. After adding 600 µl ethanol, the resulting mixture was loaded onto a DNeasy Spin column following manufacture’s protocol, with a final elution volume of 100 µl. The DNA was further processed with a Zymo Research One Step PCR Inhibitor Removal kit (Zymo Research, Irvine, California, USA). A negative control was simultaneously extracted to test for possible laboratory contamination.

#### Molecular Analysis

To analyze the DNA mixture within each collected water sample, a specific DNA region of interest is targeted and copied millions of times through a process known as PCR. For freshwater mussel analysis, each water sample was amplified for a ∼175 base pair (bp) fragment of the mitochondrial 16S gene region which has previously been designed and validated for detection of freshwater mussels from eDNA samples (Prié et al. 2021, Marshall et al. 2022). Mussel eDNA was sequenced with MiSeq Illumina metabarcoding as previously described in Marshall et al. (2022).

Each water sample was amplified for a ∼175 bp fragment of the mitochondrial 16S gene region which has previously been tested for amplification of unionid mussels from eDNA samples (Prié et al. 2021, Marshall et al. 2022). Mussel eDNA was amplified and sequenced with MiSeq Illumina metabarcoding as previously described in Marshall et al. (2022). Initial PCR amplification was completed for each sample in triplicate with 10 µl PCR reactions containing 4 µl extracted eDNA, 0.4µM primer, and Applied Biosystems™ TaqMan™ Environmental Master Mix 2.0. The amplifications started with an initial denaturation at 95°C for 5 min, followed by 35 cycles of 95°C for 15s, 5% ramp down to 55°C for 30s, and 72°C for 30s. Triplicate PCR products were diluted 1:10 then pooled prior to starting the Illumina adaptor and barcoding PCR processes.

A PCR process initiated the incorporation of Illumina adaptors and multiplexing barcodes using the initial forward and reverse primers containing 33 or 34 base pairs of 5’ Illumina hanging tails to provide a priming site for a final PCR to incorporate barcodes and remaining base pairs of Illumina adaptors. The 12 µl PCR reaction contained 2 µl diluted pooled PCR product, 0.3 µM Illumina adaptor primers and 6 µl 1X Qiagen Plus Multiplex Master Mix. The PCR process denatured for 95°C for 5 min, 5 cycles of 98°C for 20s, 1% ramp down to 65°C for 15s, and 72°C for 15s., followed by 7 cycles of 98°C for 20s, 5% ramp down to 65°C for 15s, and 72°C for 15s. PCR product was diluted 1:10 prior to use in the barcode adaptor PCR process.

The final PCR incorporated paired-end dual indices (eight base pair barcodes) that allowed samples to be identified in the raw read data, and the p5/p7 adaptor sequences to allow the sample to bind onto the Illumina MiSeq flow cell. This final 12μl PCR reaction contained 1μl diluted product from the previous PCR, 0.3 µM forward and reverse indexed primer and 6ul 1X KAPA HiFi HotStart Ready Mix PCR Kit (Roche Diagnostics, Indianapolis, Indiana, USA). Conditions were 3 minutes of initial denaturation at 95°C, followed by 10 cycles at 98°C for 20 s, 5% ramp down to 72°C for 15 s, with a final 5 min 72°C extension. All PCRs were completed on Bio-Rad C1000 Touch Thermal Cyclers. Illumina adapted PCR products were pooled with equal volumes, then size selected (target ∼319bp) using 2% agarose gel electrophoresis. The final pool was sequenced with 2× 300 nt V2 Illumina MiSeq chemistry by loading 6.4 pmol library. An additional 20% PhiX DNA spike-in control was added to improve data quality of low base pair diversity samples. Additionally, a PCR no-template negative control was run for each library preparation step.

#### Bioinformatic Processing & Taxonomic Identification

Taxonomic nomenclature follows the Freshwater Mollusk Conservation Society (FMCS 2025). The metabarcoding data was processed following a bioinformatic pipeline previously described in Marshall et al. (2022). The MiSeq runs were processed separately. The forward and reverse primer sequences were removed from the demultiplexed sequences using the cutadapt (Martin et al. 2011) plugin within QIIME 2 (Bolyen et al. 2019). Next, sequence reads were filtered and trimmed using the denoising DADA2 (Callahan et al. 2016) plugin within QIIME 2. Based on the quality scores from the forward and reverse read files, a “truncLen” was set to 120 for the forward and 110 for the reverse read files. Using DADA2, error rates were estimated, sequences were merged and dereplicated, and any erroneous or chimeric sequences were removed. Unique sequences were then clustered into Molecular Operational Taxonomic Units (MOTUs) using the QIIME 2 vsearch de-novo with a 97.5% similarity threshold (Coghlan et al. 2021, Marshall et al. 2022). MOTUs from unionid taxa were identified to the species-level using the Basic Local Alignment Search Tool (BLAST+, https://blast.ncbi.nlm.nih.gov/Blast.cgi; Camacho et al. 2009) against our custom database of both in-lab generated sequences and mt-16S sequences downloaded from NCBI GenBank. These MOTUs were further validated with comparisons against the complete NCBI nr database, to investigate alignment to mis-labeled sequences or species not historically within the sampling region.

Freshwater mussels display a unique form of mitochondrial inheritance, termed doubly uniparental inheritance (DUI), in which males possess a paternal mitochondrial mitotype that is largely restricted to male gonads and gametes (Gusman et al. 2016). As the male mitotype is genetically distinct from the female mitotype (Curole & Kocher 2005), detections originating from the female or male mitotype were separated within the metabarcoding dataset. The male lineage for *S. ambigua* was able to be detected in the current study because a voucher swab was collected and used to generate the first reference sequence, whereas a majority of species lack reference genetic sequences for their male lineage and cannot currently be detected within an eDNA dataset (Marshall et al. 2025, Marshall et al. 2026a).

Sequences were assigned to a species if they met a threshold of >97.5% identity and 100% query coverage. Furthermore, sequences that were assigned to multiple species with the same BLAST e-value score were inspected and a final decision was made based on known distribution and presence within the sampled watershed. Additionally, if multiple sequences were assigned to the same taxonomy, they were inspected and removed or collapsed into a single MOTU to obtain a final matrix of read counts per taxa.

In metabarcoding analysis, the sequencing process can introduce a form of sample cross-contamination in which sequences from one sample are falsely detected within another sample, often termed ‘critical mistags’, ‘tag jumps’, or ‘index hopping’ (Esling et al. 2015; Bohmann et al. 2022, Richardson 2022).

These sequencing artifacts generally represent a small fraction of the total sequences (Esling et al. 2015; Schnell et al. 2015), yet they can lead to false detections in some circumstances. Therefore, the final processing of the MOTUs in this study consisted of estimating and removing mistags from the dataset following the framework outlined by Richardson (2022). Based on the number of reads per MOTU within the field and laboratory negative controls, the mistag rate was estimated for all dual-index combinations. The estimated mistag rate of the current dataset was 0.001, but a conservative estimate of 0.0075 was used for identifying and removing mistags (Richardson 2022). Any sequences identified as field contamination were not included in estimating the mistag rate (see Results).

### Statistical Analysis

To assess the repeatability of eDNA detection, an eDNA Detection Index was developed and applied to all species detections at each site. The index levels are described as:

- *Non-detection* occurs when a species is not detected in any water sample replicates collected within a single site.
- *Low repeatability* occurs when a species is detected in < 50% of water replicates collected within a single site.
- *Moderate repeatability* occurs when a species is detected in < 75% and ≥ 50% of water replicates collected within a single site.
- *High repeatability* occurs when a species is detected in ≥ 75% of water replicates collected within a single site.

#### Comparing Visual and eDNA Surveys

The total observed mussel abundance and the total eDNA sequence read counts across all of Fish Creek were calculated for each mussel species detected. A bubble plot and bar plot were generated to visualize the detected mussel assemblage from the two survey methods.

##### Sampling Site Level Detections

Because the amount of survey effort varied across sites (i.e., the amount of search time), the total observed mussel abundance for each species at a site was standardized by calculating the Catch Per Unit Effort (CPUE), a measure of the mussel abundance recovered per hour of search. Similarly, the site specific eDNA sequence read counts were calculated by averaging the total read counts across replicate water samples collected within a sampling site.

Bubble and bar plots were generated in R to visualize the CPUE from visual surveys and the mean eDNA read sequence count within each sampling site for each detected species.

The total species detections from eDNA surveys were compared to the total visual observations. The proportion of eDNA detections occurring at a site with confirmed visual presence were calculated across the repeatability scale of eDNA detection. Furthermore, logistic regression was used to assess the repeatability of eDNA detection based on the observed species abundance within each sampling site.

The mean eDNA read sequence counts and the observed mussel abundance for each detected mussel species within each of the 28 visual surveyed sites was compared using a logarithmic regression. The number of observed CPUE per site and the number of eDNA sequence counts per site for each detected species within each of the 28 visual surveyed sites were assessed based on the repeatability scale of eDNA detection using a Kruskal–Wallis test and visualized using boxplots.

#### Evaluating eDNA Survey Protocol

The proportion of occupied sites for each mussel species was compared between the two survey methods using logistic regression. The proportion of occupied sites for each mussel species from the eDNA survey was calculated for each of the three levels of repeatability of eDNA detection. The proportion of occupied sites for each mussel species from the visual surveys and the eDNA surveys was compared to the total observed mussel abundance using logistic regression. The proportion of occupied sites for each mussel species from the eDNA survey was calculated for each of the three levels of repeatability of eDNA detection. The eDNA detection of a species at a site was compared to the observed CPUE at that site, with eDNA detections grouped by the detection repeatability scale. This comparison included a logistic regression to examine the influence of observed CPUE on the likelihood of eDNA detection.

To assess the ability of eDNA to estimate stream wide occupancy, Bayesian integrated occupancy models were developed in the ‘spOccupancy’ R package (Doser et al. 2022). To accomplish this, the eDNA relative-abundance data was transformed into binary detection/non-detection data for all species detected with eDNA across Fish Creek. Occupancy models leverage repeated observations of detection/non-detection data to jointly estimate detectability and probability of occurrence (MacKenzie et al., 2002). This multispecies single-season occupancy model was evaluated for all species to estimate the mean occupancy (ψ) (i.e., the estimated number of occupied sites) and the mean detection probability (*p*) (i.e., the probability of successful eDNA detection of a species within a replicate environmental sample). We hypothesized that species would vary in their probability of occupancy throughout Fish Creek based on (1) drainage area and (2) land use and riparian buffer conditions. To account for these factors, we included distance from the St. Joseph River confluence as a site-level covariate, as this distance likely serves as a proxy for both drainage size and habitat characteristics along Fish Creek.

Next, the estimated probability of eDNA detection for each species was compared to their mean observed CPUE during the visual surveys. Finally, the survey design was evaluated by calculating the detection probability (*p*\*) for each species: *p*\* = 1-(1-*p*)^n^ where *p* is the estimated detection for a single replicate environmental sample and n is the total number of replicates. The survey effort was estimated to exceed 0.75 and 0.95 detection probabilities for each species.

## Results

### Bioinformatic Processing

The raw MiSeq output included a total of 27,205,367 sequences, of which 14,547,447 reads were retained and identified as freshwater mussel.

One of the field control samples (eDNA Site FC22-4 [sample FC22-4G]) resulted in high detection of mussel sequences (16,060 sequence reads) and high number of species (15 MOTUs). This field control was extracted in the same batch as replicate A from eDNA Site FC22-5 (sample FC22-5A), which subsequently resulted in no mussel reads. It is possible that an inadvertent swap of samples occurred between these two during the sample shipment and/or laboratory processing. The field control sample from eDNA Site FC22-4 provided detections for species that were likewise found at Site FC22-5.

However, we cannot definitively conclude if this occurred. We therefore remove samples FC22-4G and FC22-5A from the analyzed dataset. We include the data from the other replicate samples at FC22-4 and FC22-5 in the analysis below, but note that these sites may have been exposed to cross -contamination.

Of the remaining 21 field and laboratory controls, the number of mussel DNA sequences ranged from 0 to 632 per control sample. For the MOTU sequences found within the control samples, their sequence count ranged from 8 to 686 total reads. The MOTUs found within the control samples pertained to the most common MOTUs within the dataset. Following the mistag analysis, 2% (32 of 1614) of the detections from the eDNA samples were classified as being within the range of probable mistag, and thus they were removed.

### Taxonomic Identification

Of the species historically found within Fish Creek (Watters 2000), two species are currently lacking sequence data for the 16S gene region within available genetic databases – *E. perobliqua* and *Toxolasma lividum* (Purple Lilliput). While no genetic data are available for *E. perobliqua*, a previous genetic voucher was collected for the closely related *Epioblasma obliquata* (Purple Cat’s Paw Pearly Mussel) (Marshall et al. 2022), which was used as a reference to identify any closely related sequences within the eDNA data. The metabarcoding data from Fish Creek resulted in 25 taxa assigned as a unionid female mitotype spanning across five unionid tribes (Table 1, Supplementary Figure 1).

These taxa ranged from 97.8-100% identity to a species within the curated genetic database. Twenty of the 25 were exact matches to a reference genetic sequence, while only *Cyclonaias tuberculata* (Purple Wartyback) and *Theliderma cylindrica* (Rabbitsfoot) were below 99% identity. While there are no genetic reference sequences for the 16S gene for Purple Lilliput, it is unlikely this species was detected and unidentified in the current study. Across other mitochondrial gene regions (COI and ND1), Purple Lilliput and *Toxolasma parvum* (Lilliput) display >10% divergence, suggesting these two species would also be easily distinguishable with the 16S gene region used in this study.

### Stream-wide Species Assemblage

Visual surveys found 22 live mussel species across the sampling region of Fish Creek (Table 1, Supplementary Figure 2). The eDNA survey detected 25 total species, including 21 of the 22 found during the visual survey (Table 1, Supplementary Figure 2). *Ortmanniana ligamentina* (Mucket) was the lone species missed with eDNA, and was found as only a single individual from two sites with the visual survey (Table 1, Figure 2). The eDNA survey detected the three federally protected species found with the visual survey, which included *Paetulunio fabalis* (Rayed Bean), *Pleurobema clava* (Clubshell), and *T. cylindrica* (Table 1, Supplementary Figure 2). The eDNA survey detected four species that were not found during the visual survey, which included *Lampsilis fasciola* (Wavyrayed Lampmussel)*, Simpsonaias ambigua* (Salamander Mussel)*, T. parvum,* and *Utterbackia imbecillis* (Paper Pondshell) (Table 1, Supplementary Figure 2). However, the visual survey did recover weathered shells of *L. fasciola* and a fresh dead shell of *S. ambigua* (Table 1).

**Figure 2.**
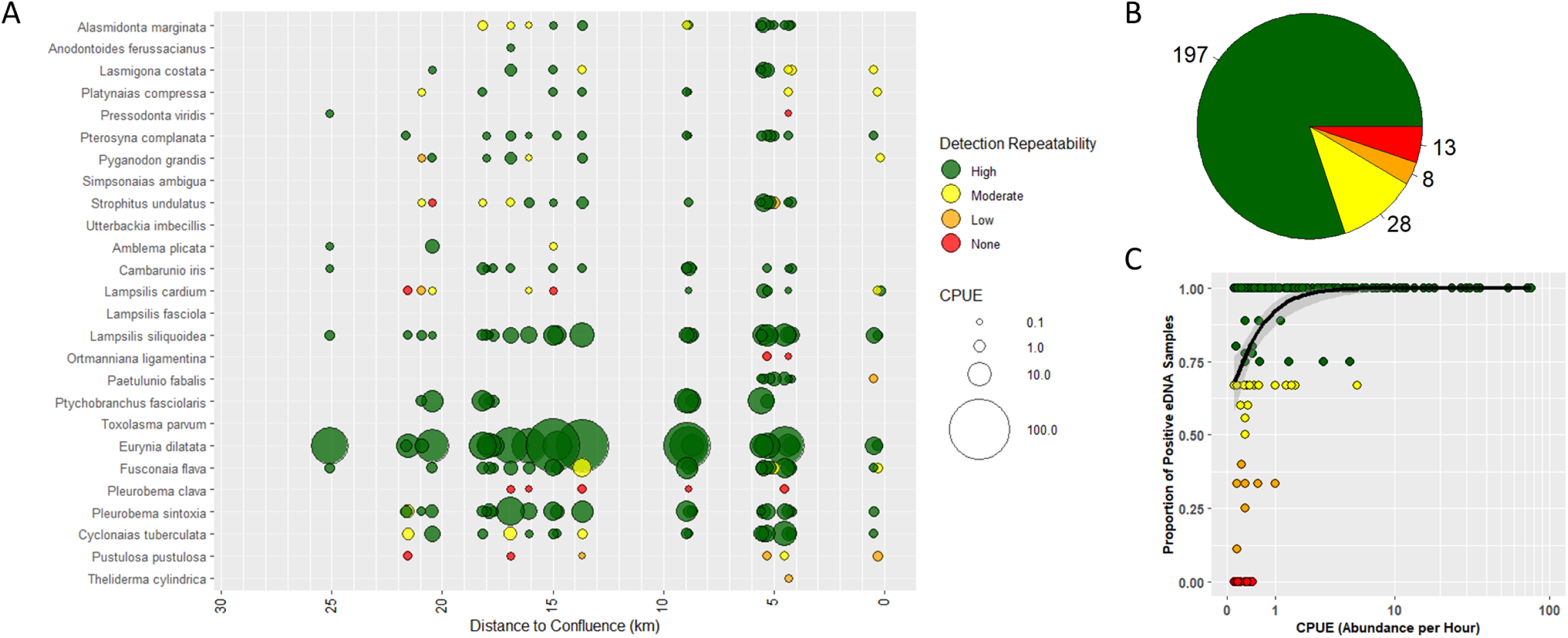
(A) Bubble plot of visual observations for each species across Fish Creek. Colors represent eDNA detection repeatability for all visual observation. (B) Pie chart of the number of occurrences for each eDNA detection repeatability for visual observations. (C) Logarithmic regression of the proportion of eDNA detections for the catch-per-unit-effort of each visual observation.

Visual surveys recovered a total of 4,326 mussels, with abundances ranging from one for *Anodontoides ferussacianus* (Cylindrical Papershell) to 2,702 for *Eurynia dilatata* (Spike) (Table 1, Supplementary Figure 2). The eDNA surveys resulted in a total of 14,547,041 female DNA sequence reads of freshwater mussels, with species sequence counts ranging from 842 for *T. parvum* to 7,425,092 for *E. dilatata* (Table 1, Supplementary Figure 2). The four species that were detected with eDNA only (*L. fasciola, S. ambigua, T. parvum,* and *U. imbecillis*) accounted for less than 0.2% of the total eDNA sequence counts (Table 1, Supplementary Figure 2). *Simpsonaias ambigua* was later confirmed present at river kilometer 5.55 after intensive searching that tripled the level of visual search effort (Marshall et al. 2026b). Estimates of mussel composition across Fish Creek from visual survey abundances were positively correlated with eDNA sequence counts (R^2^ = 0.61, *p* < 0.001***; Supplementary Figure 2).

### Site-level Species Assemblages

Across Fish Creek, both surveys found mussel assemblages were dominated by *E. dilatata, Lampsilis siliquoidea* (Wavy Rayed Lampmussel), and *Ptychobranchus fasciolaris* (Kidneyshell) (Figure 2, Supplementary Figure 3, Supplementary Figure 4). Of the sites that were surveyed with both visual and eDNA surveys, visual surveys observed a species at a site on 246 occasions, while eDNA observed a species at a site on 490 occasions (Figure 2A; Supplementary Figure 3). Of the visual observations, eDNA provided a corresponding positive detection on 233 (94.7%) occasions (Figure 2B). In most instances, if a species was visually observed it was likewise detected with eDNA at high repeatability (197 of 246 occasions [80.1%]), whereas visual observations were less often detected as moderate or low repeatability (moderate: 23 of 246 occasions [11.4%], low: 7 of 246 occasions [3.3%]) (Figure 2B). Twelve of the 13 (92.3%) times a species was missed at a site with eDNA the visual survey observed only a single individual for that species. Furthermore, as the CPUE increased for an observed species at a site, so did the proportion of positive eDNA sample replicates (Figure 2C).

Of the eDNA observations across these 28 sites, 334 occurred as high repeatability detections (Supplementary Figure 3). Of these high repeatability detections, 137 (41.0%) occurred at a site where a species that was absent in the visual survey. Whereas the detections for moderate repeatability and low repeatability had a greater number of observations not coinciding with a visual observation (moderate: 56 of 84 [67.7%], low: 64 of 72 [88.9%]) (Figure 3).

**Figure 3.**
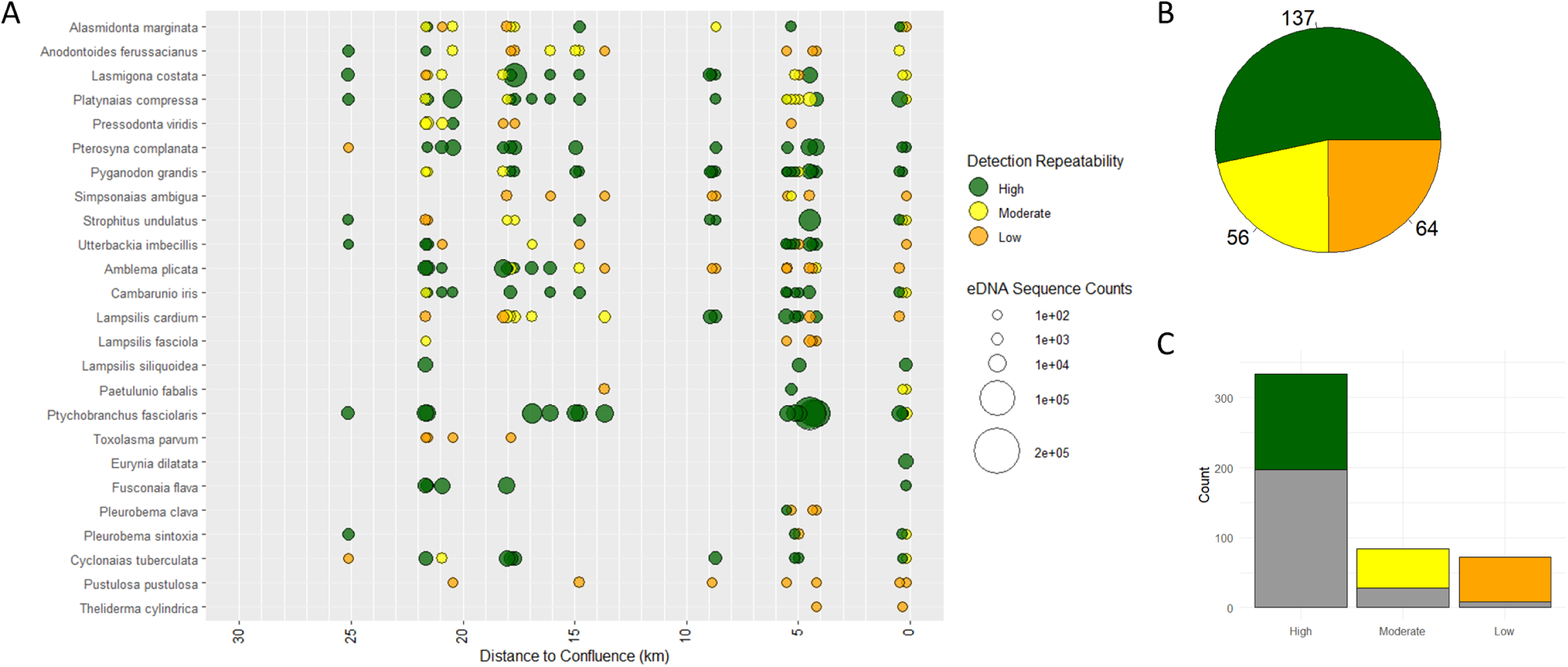
(A) Bubble plot of eDNA detections occurring without a visual observation for each species across Fish Creek. Colors represent eDNA detection repeatability for all eDNA detections occurring without a visual observation. (B) Pie chart of the number of occurrences for each eDNA detection repeatability for all eDNA detections occurring without a visual observation. (C) Barchart of all eDNA detections based on eDNA detection repeatability. Grey bar represent eDNA detections that coincide with a visual observation.

Across the sites with visual and eDNA surveys, the mean eDNA sequence count per site was positively correlated with the mean CPUE for each species (R^2^ = 0.63, *p* < 0.001***) (Figure 4A). When a species was detected with eDNA at high repeatability, its eDNA was abundant within the environment, and the detection corresponded to species that were observed at higher CPUE. For example, across the eDNA detection repeatability scale, the observed CPUE was greatest for eDNA detections of high repeatability (Figure 4B). Similarly, the eDNA sequence count was greatest for detections of high repeatability (Figure 4C).

**Figure 4.**
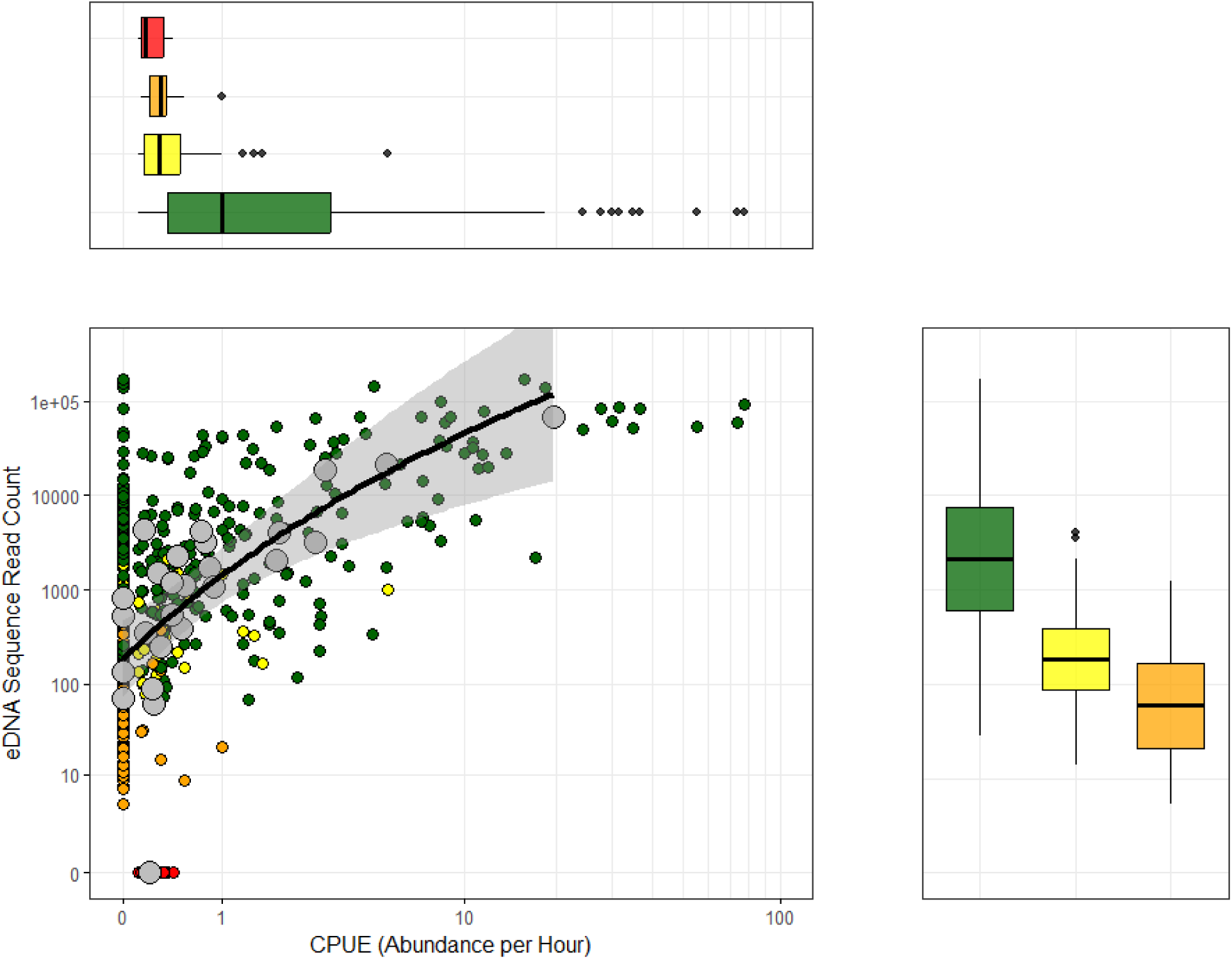
(A) Logarithmic regression of eDNA sequence count compared to visual mussel abundance across the sites with both eDNA and visual surveys. Large grey points indicate the mean eDNA sequence count or visual abundance for each mussel species, small circles represent site-level detections for each mussel species. (B) Boxplot of mussel abundance from visual surveys across the repeatability scale of eDNA detection. (C) Boxplot of eDNA sequence count from eDNA surveys across the repeatability scale of eDNA detections.

#### Environmental DNA Detection Efficiency

The median probabilities of eDNA detection ranged from 0.079 (0.016 – 0.389) for *T. parvum* to 0.995 (0.974 – 0.999) for *L. siliquoidea* and *E. dilatata* (Figure 5A). Most species (19 of 25) had baseline detection probabilities greater than 0.50 (Figure 5A). The median probability of detection estimated across mussel species were positively correlated with the mean CPUE observed during the visual surveys (R^2^ = 0.56, *p* < 0.001***) (Figure 5B). The analysis of sampling effort (i.e., number of eDNA samples per site) required to reach probability of detection threshold suggests the rarest observed CPUE of 0.16 would surpass a 0.75 probability of eDNA detection at four samples (2 – 12 CI) per site (Figure 5C) and surpass a 0.95 probability of eDNA detection at nine samples (4 – 26 CI) (Supplementary Figure 5).

**Figure 5.**
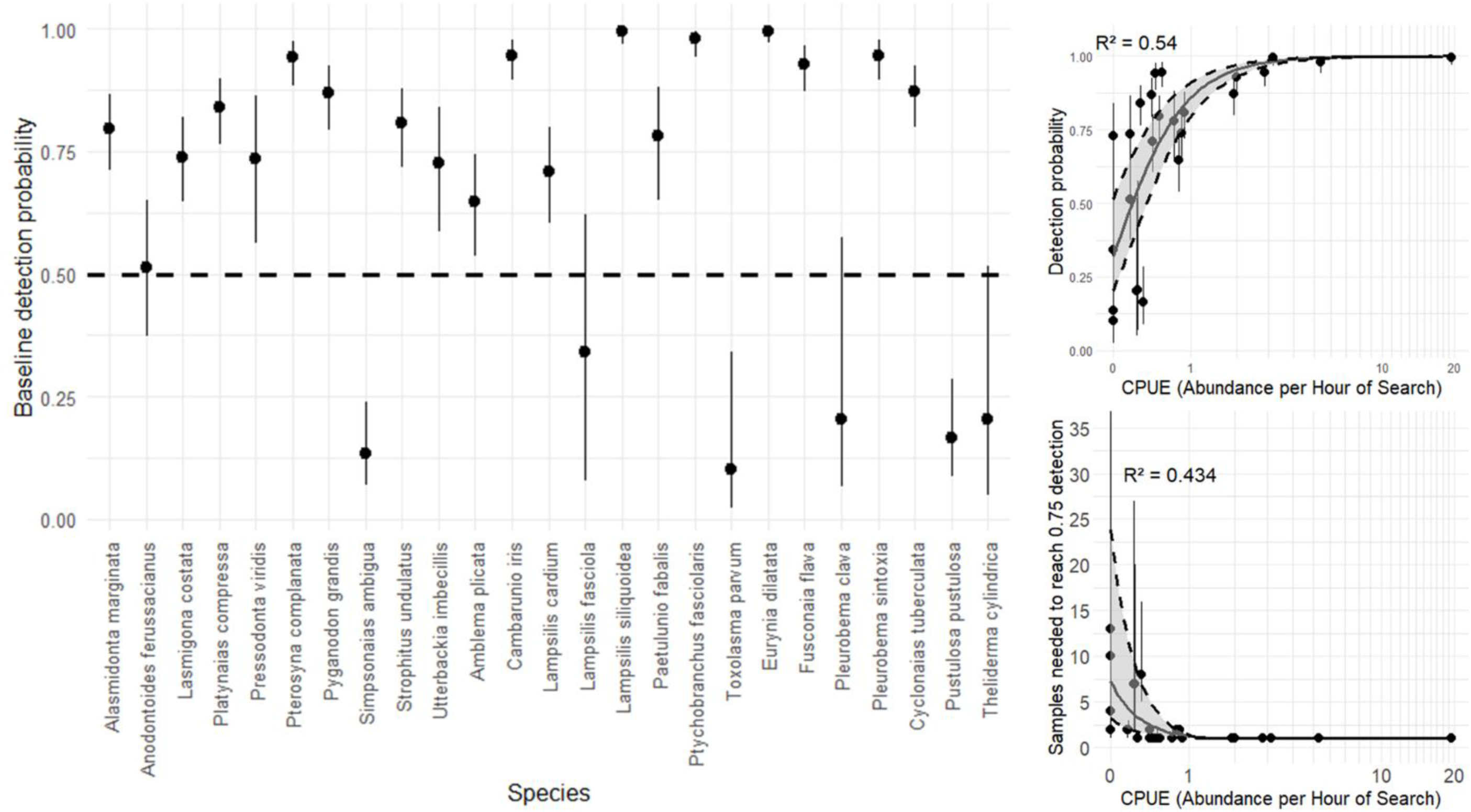
(A) The estimated median baseline environmental DNA detection probability for each mussel species detected. (B) The estimated median baseline eDNA detection probability compared to the observed catch-per-unit-effort (CPUE) for each species. (C) The estimated number of samples required to exceed a 0.75 probability of detection compared to the observed CPUE for each species.

### Site Occupancy Estimates

Of the 28 sites surveyed with visual surveys, the most widespread species were *E. dilatata* (27 sites), *L. siliquoidea* (25 sites), *Fusconaia flava* (Wabash Pigtoe) (23 sites), and *Pleurobema sintoxia* (Round Pigtoe) (23 sites) (Figure 2). These four species were likewise widespread in the eDNA survey, detected at all 32 eDNA sampling sites (Supplementary Figure 3). In addition, three other species were detected with eDNA at all 32 sampling sites, *Cambarunio iris* (Rainbow), *Platynaias compressa* (Creek Heelsplitter), and *P. fasciolaris* (Supplementary Figure 3).

The proportion of occupied sites for a species positively increased with observed CPUE within the visual and eDNA surveys (Supplementary Figure 6A), however, the modeled curve for eDNA data best resembled the visual survey when using eDNA detections of high repeatability only (Supplementary Figure 6A). Similarly, comparisons of proportion of occupied sites between survey methods best approximated a 1:1 relationship when using eDNA detections of high repeatability only (Supplementary Figure 6B).

The occupancy model incorporating distance to the St. Joseph River confluence as a site-level covariate performed better than the null model (WAIC of 1876.82 compared to 2024.57 for the null). There was strong congruence between the estimated occupancy probability models based on eDNA detections in comparison to distributions of the visual observed occupancy (Figure 6). Across the sites with visual observations for all species within Fish Creek, 90% of these species occurrences (220 of 244) aligned with an estimated site occupancy of greater than 0.95 from the eDNA occupancy models, and the mean estimated occupancy probability across all visual occurrence points was 0.96 (Figure 6 and Supplementary Figure 7). Multiple species that were visually observed to be stream-wide were estimated to have high occupancy across the entire stream with eDNA (Figure 6). One exception to the congruency between eDNA occupancy and visual observations occurred for *P. clava*, which was observed at five sites during visual surveys (spanning from four to 17 kilometers from the confluence), but only detected within four to five kilometers from the confluence with eDNA (Figure 6).

**Figure 6.**
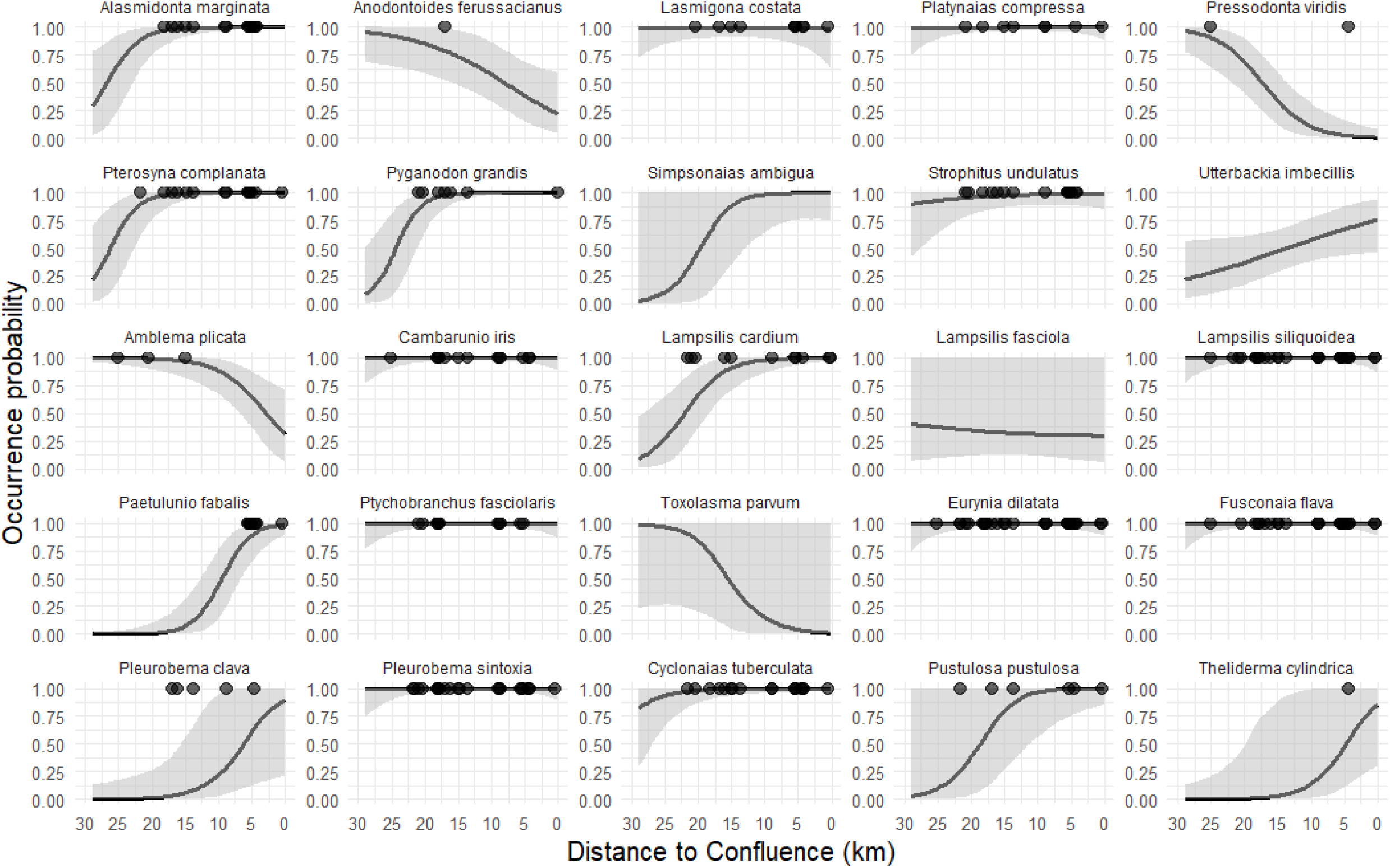
The estimated occupancy across Fish Creek for each species based on environmental DNA sampling. The black points indicate sites with visual observations.

The estimated probability of occupancy was negatively influenced by distance from the confluence with the St. Joseph River for seven of the 25 species (Figure 6 and Supplementary Figure 8), whereas three species were positively influenced by distance from the confluence and were more likely to be detected in the headwaters of Fish Creek (Figure 6 and Supplementary Figure 8). For example, *Amblema plicata* (Threeridge) and *Pressodonta viridis* (Slippershell) had greater eDNA sequence counts and greater occupancy estimates for sites greater than 15 kilometers from the confluence, coinciding with its visual observations (Figure 6 and Supplementary Figure 9). Conversely, *P. fabalis* and *T. cylindrica* had greater eDNA sequence counts and greater occupancy estimates for sites near the confluence with the St. Joseph River, again coinciding with visual observations (Figure 6 and Supplementary Figure 9). Distance to the confluence likely served as a proxy for underlying environmental gradients, such as land use, habitat conditions, and drainage area, and therefore should be interpreted cautiously, as these correlated factors may be the true drivers of the observed patterns.

## Discussion

This study used eDNA metabarcoding to characterize freshwater mussel assemblages across 30 km of Fish Creek and demonstrated that eDNA provides a broader and more sensitive assessment of species richness than the currently implemented visual surveys. The eDNA survey detected 25 species compared to the 22 species observed visually. Notably, eDNA detected four species which had no live observations during the visual surveys, but were historically known to be present within the system (Watters 2000). Both surveys confirmed extant populations of three federally protected mussel species in Fish Creek, while eDNA was the lone survey method to provide evidence for the presence of the proposed federally endangered *S. ambigua*. This species is particularly hard to find with conventional surveys due to behavioral characteristics and habitat use, in which it is often buried in the stream bank, bedrock crevices, or under large flat boulders (Porto-Hannes et al. 2021). Additionally, previous eDNA surveys have demonstrated increased sensitivity for *S. ambigua*, especially at locations with low abundance (Douglas et al. 2025, Marshall et al. 2026b). The lack of detection for *E. perobliqua* from either visual or eDNA surveys may suggest its true extirpation from the system.

Across sites, 95% of visual observations were matched by eDNA, indicating that the eDNA survey provided reliable detection for species that were truly present. However, more than half of all eDNA detections occurred at sites where a species was not visually observed, and eDNA generally detected species at more sites than visual surveys. These patterns likely reflect some combination of missed detections due to the relatively low effort visual searches and downstream transport of eDNA. Hydrological processes can move eDNA over long distances (Wilcox et al. 2016; Van Driessche et al. 2022), and previous studies have documented bivalve eDNA transport exceeding 1 km in flowing systems (Deiner & Altermatt 2014; Shogren et al. 2019; Stoeckle et al. 2021). Because visual surveys sampled only short (∼25 m) riffle sections, whereas eDNA integrates information across a larger hydrological footprint, the two methods inherently represent different spatial scales of detection. Even though eDNA provided a greater number of overall detections, the occupancy models based on eDNA displayed high congruence in occupancy estimates to locations of ground-truthed visual observations.

Differences in habitat associations may further explain mismatches between methods. For example, *U. imbecillis* and *T. parvum* prefer soft substrates and slackwater habitats (Watters et al. 2009). These two species were not visually observed within Fish Creek but were both detected with eDNA, including *U. imbecillis* being detected from half of the sites. Visual surveys targeted riffles where diverse mussel assemblages typically occur, but these habitats are not optimal for all species. Thus, eDNA likely captured upstream or adjacent assemblages not represented in the extent of the visually surveyed habitat.

Using a scale of eDNA detection repeatability provided better congruence with visual observations. For example, the majority of eDNA detections at high repeatability corresponded with a visual observation, whereas the majority of moderate to low repeatability did not correspond with visual observations. High repeatability detections were strongly associated with visual presence, whereas moderate and low repeatability detections were less likely to correspond to site-level occupancy. This suggests that repeated detection across sample replicates is a useful indicator of local presence, while lower repeatability may reflect broader reach scale occupancy corresponding to eDNA transport.

Species abundance strongly influenced eDNA detection probability, consistent with previous studies (Marshall et al. 2022, Marshall & Fleece 2025, Marshall et al. 2026c). Similarly, eDNA counts were correlated with the observed mussel CPUE at a site. These correlations indicate that metabarcoding can provide a coarse index of mussel abundance despite known limitations of metabarcoding related to species-specific shedding rates, biomass differences, and primer biases (Ruppert et al. 2019, Yates et al. 2021). Even with substantial variation in size (e.g., *P. fabalis* adults reach ∼40 mm, whereas *A. plicata* can reach ∼170 mm; Watters et al. 2009) and diverse life history traits across mussel tribes (Haag 2012), eDNA reliably distinguished abundant, common, and rare taxa across Fish Creek.

The low effort eDNA survey proved sufficient for detecting rare species. For example, the lowest observed CPUE of 0.16 was estimated to exceed a 0.75 detection probability with four eDNA samples. However, achieving a 0.95 detection probability required nine samples, and thus greater sampling rigor is required for site-specific assessments compared to watershed-level assessments. These estimates apply to a small, low flow stream and larger systems will likely require greater sampling effort, as was seen in the Walhonding River, Ohio (Marshall & Fleece 2025). Field efficiency was also substantially higher for eDNA: three replicate water samples required roughly 40 minutes per site, compared to an average of 4.5 hours for visual surveys. However, eDNA metabarcoding involves laboratory processing which may require weeks to months to produce results, whereas visual surveys provide immediate, actionable information. Thus, the choice between methods should consider project goals, timelines, and available resources.

While rare species were routinely detected with eDNA, three federally protected species (*P. clava, T. cylindrica*, and *S. ambigua*) were never detected above low to moderate repeatability, even at sites where *P. clava* and *T. cylindrica* were visually observed. This demonstrates that rare species may have inconsistent eDNA detection and require greater sampling effort. For *P. clava*, eDNA was detected at four sites between 4 to 6 kilometers from the confluence, whereas visual surveys observed *P. clava* up to 17 kilometers from the confluence.

The lower detection of *P. clava* with eDNA may be a result of its rarity within the mussel community and its life history traits. From the sites it was present at had the highest overall mussel abundances, and *P. clava* always represented less than 0.5% of the total mussel community. Rare species’ eDNA fragments are less likely to be captured and sequenced as a result of primer-bias and competition during PCR amplification (McClenaghan et al. 2020). Additionally, *P. clava* buries deeply in the substrate (Smith et al. 2001, Watters et al. 2009), potentially reducing eDNA release into the water column. Seasonal variation in eDNA detection has been documented for some endobenthic mussels (Marshall et al. 2026d, Johnson et al. 2025), and the low detection may reflect seasonal endobenthic behavior for *P. clava*.

When implications on species presence at a site-level are high, greater sampling rigor will increase confidence for probable absence determinations in cases of non-detections. Although single-species molecular assays such as qPCR and ddPCR are often assumed to be more sensitive than metabarcoding for detecting rare taxa (Bylemans et al. 2019), this pattern is not universal. For example, no significant improvement in detecting *S. ambigua* was observed when comparing the commonly used Prié et al. (2021) mussel metabarcoding assay with a species-specific qPCR assay (Marshall et al. 2026b). The strong performance of the metabarcoding assay likely reflects its relatively narrow taxonomic specificity, targeting only unionid mussels, because metabarcoding sensitivity is strongly influenced by how broad or restricted the assay’s taxonomic scope is (Ficetola et al. 2015, Deiner et al. 2017). This illustrates that metabarcoding can effectively detect rare DNA targets when the assay’s specificity is sufficiently constrained.

## Conclusion

This study demonstrated that eDNA, if used alone, can produce data that is directly applicable for conservation planning and recovery efforts: (1) high probability of species detection with a relatively small sample size; (2) eDNA sequence counts correlate with relative abundances; and (3) eDNA detections provide species distributions. However, molecular techniques are even more powerful when they are used to compliment, not replace, visual tactile methods. In some respects, the larger “source area” sampled via eDNA may be advantageous for detecting rare species. For example, a phased sampling approach where eDNA is collected first and followed by intensive surveys may be an effective way to refine the range of especially rare species. In this case, *S. ambigua* follows that example, where it was first detected with eDNA and later verified (Marshall et al. 2026b). Additionally, metabarcoding methodology provides opportunities to simultaneously assess multi-species of conservation concern. For example, while the primary conservation target of the current study was *E. perobliqua*, the confirmation of an extant population of *S. ambigua* was an ancillary benefit of tremendous value. Together, these findings illustrate that eDNA metabarcoding can substantially enhance mussel conservation efforts when applied with an understanding of hydrological context, species ecology, and appropriate sampling rigor.

## Acknowledgments

This project was supported through field support during mussel surveys from Cassie Hauswald, William L. Chadderton, Larry Clemens, and Andrew Tucker from The Nature Conservancy, Keith Lott, Jeromy Applegate, Jennifer Finfera, Marissa Reed, Robin McWilliams, William Tucker, Lindsey Korfel, Susan Cooper, Scott Glassmeyer, Jocelynn Samu-Pittard, Liz McCloskey, Abby Bourne, and Amber Bellamy from the United States Fish and Wildlife Service, Brant Fisher and Jake Adams from the Indiana Department of Natural Resources, Megan Michael and Brittany Muncy from the Ohio Department of Transportation, and Sarah Tebbe, Kyla Maunz, Kristin Stanford, Erin Hazelton, Abby Ditomassi, Ethan Bingham, and John Navarro from the Ohio Department of Natural Resources. Additional support was provided by Roger Miller and Karen Hallberg, United States Fish and Wildlife Service.

## Data Availability Statement

Supplementary data files are provided to Open Science Framework at https://doi.org/XXXX. Raw sequence data are deposited in the National Center for Biotechnology Information (NCBI) Sequence Read Archive (SRA) under the BioProject accession XXXXX.

## Ethics and Permit Approval Statement

Tactile mussel surveys were conducted with permitted malacologists Jeromy Applegate, Keith Lott, and Megan Seymour under scientific collection permit # SC210020 and USFWS permit # 21-05.

## Permission to Reproduce Material from other sources

All material is original to this manuscript.

---

**Supplementary Figure 1.**
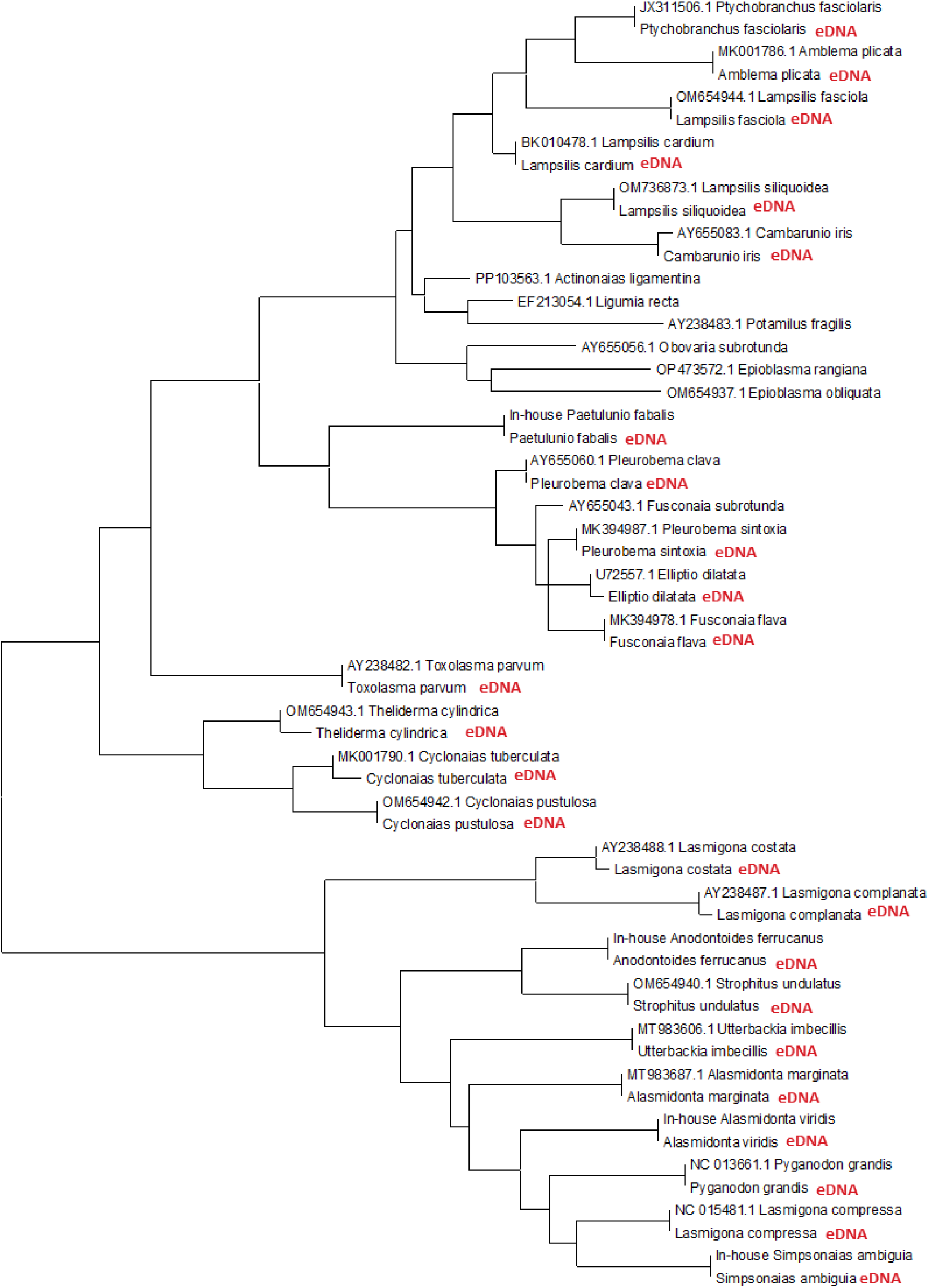
Phylogenetic tree used for taxonomic identification of the female mussel environmental DNA sequences based on the genetic match to known genetic data within the NCBI Genbank repository.

**Supplementary Figure 2.**
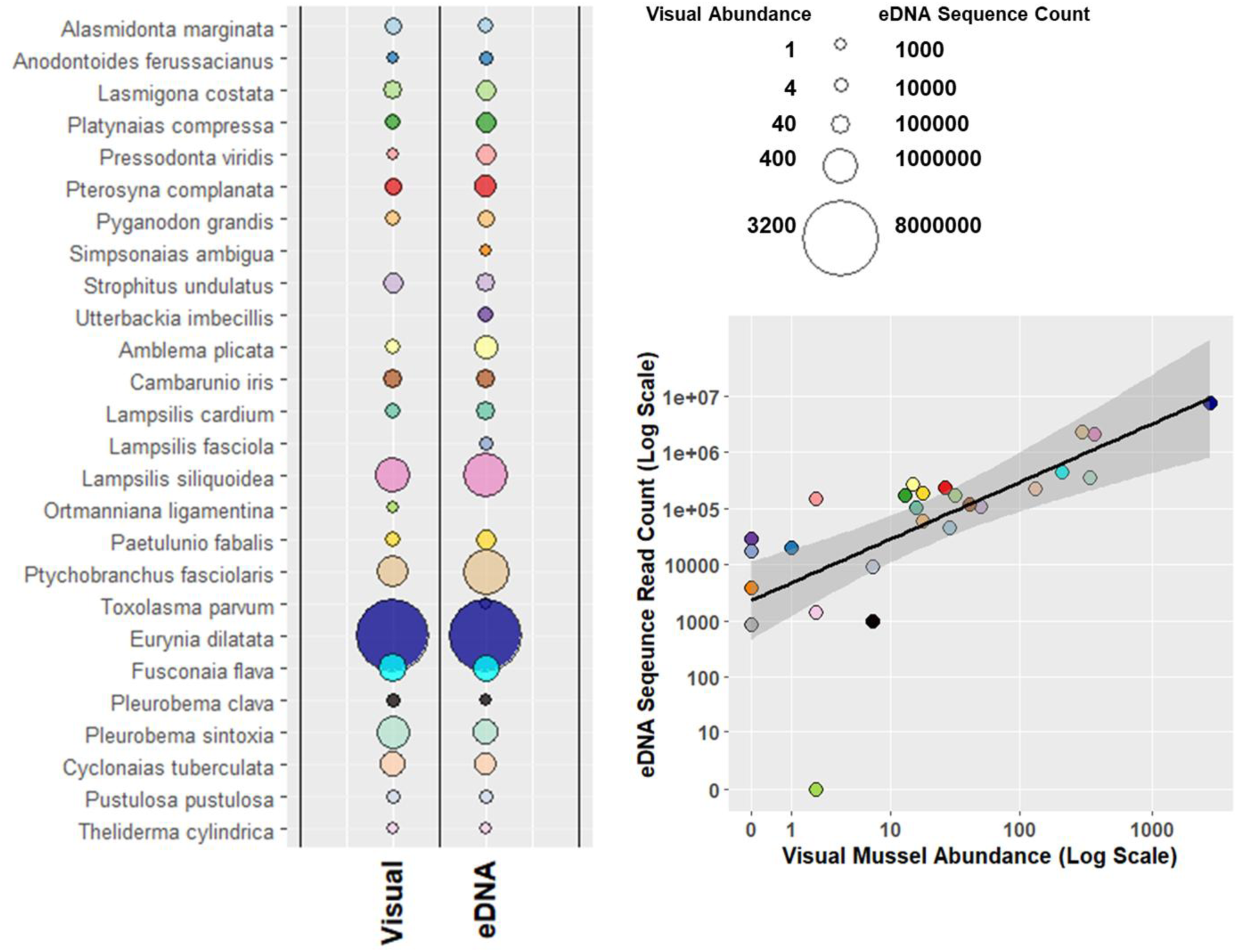
(A) Bubble plot of visual and environmental DNA (eDNA) counts across all sampling sites, and (B) linear regression of observed visual abundance to eDNA sequence counts for each detected species.

**Supplementary Figure 3.**
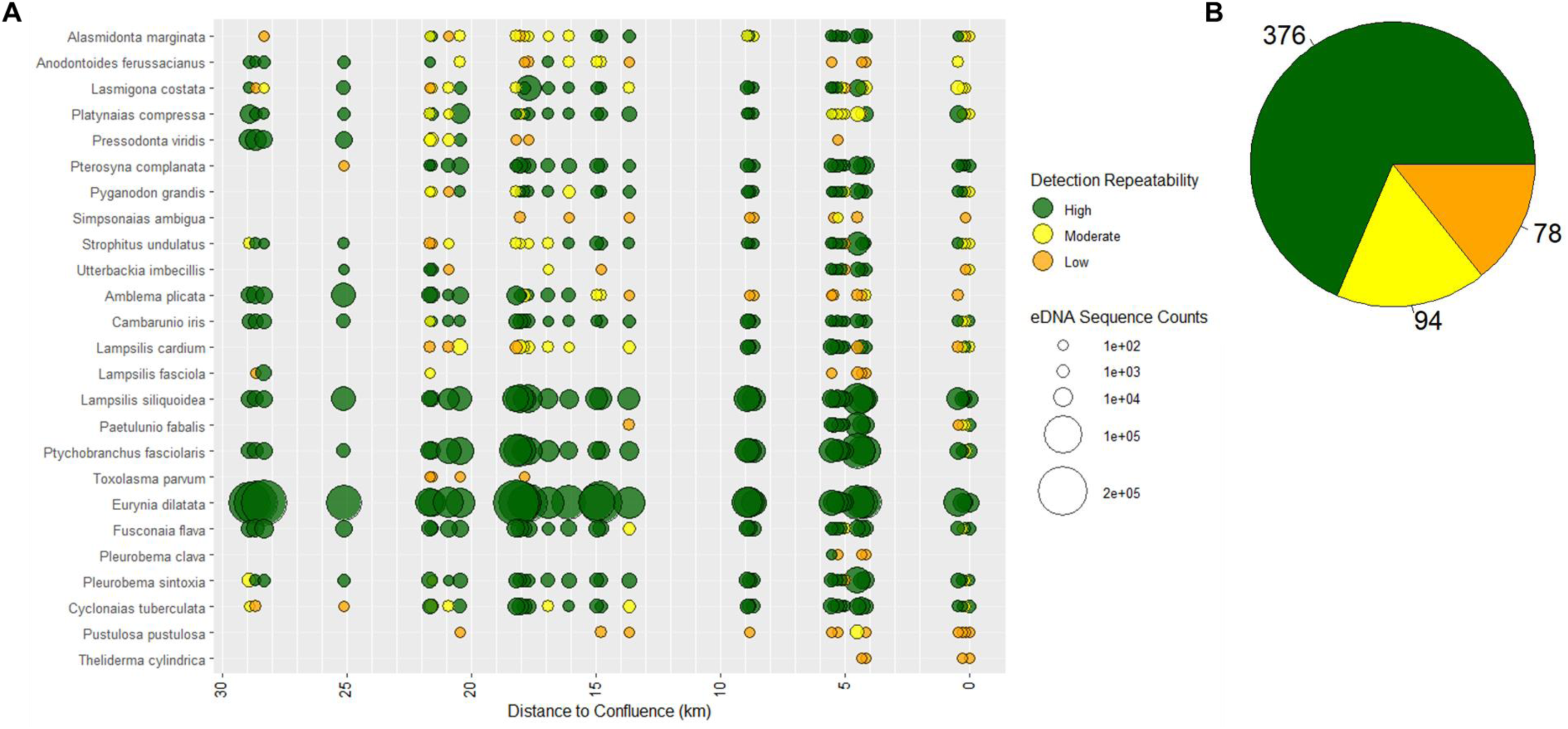
(A) Bubble plot of eDNA detections for each species across Fish Creek. Colors represent eDNA detection repeatability. (B) Pie chart of the number of occurrences for each eDNA detection repeatability for all eDNA detections.

**Supplementary Figure 4.**
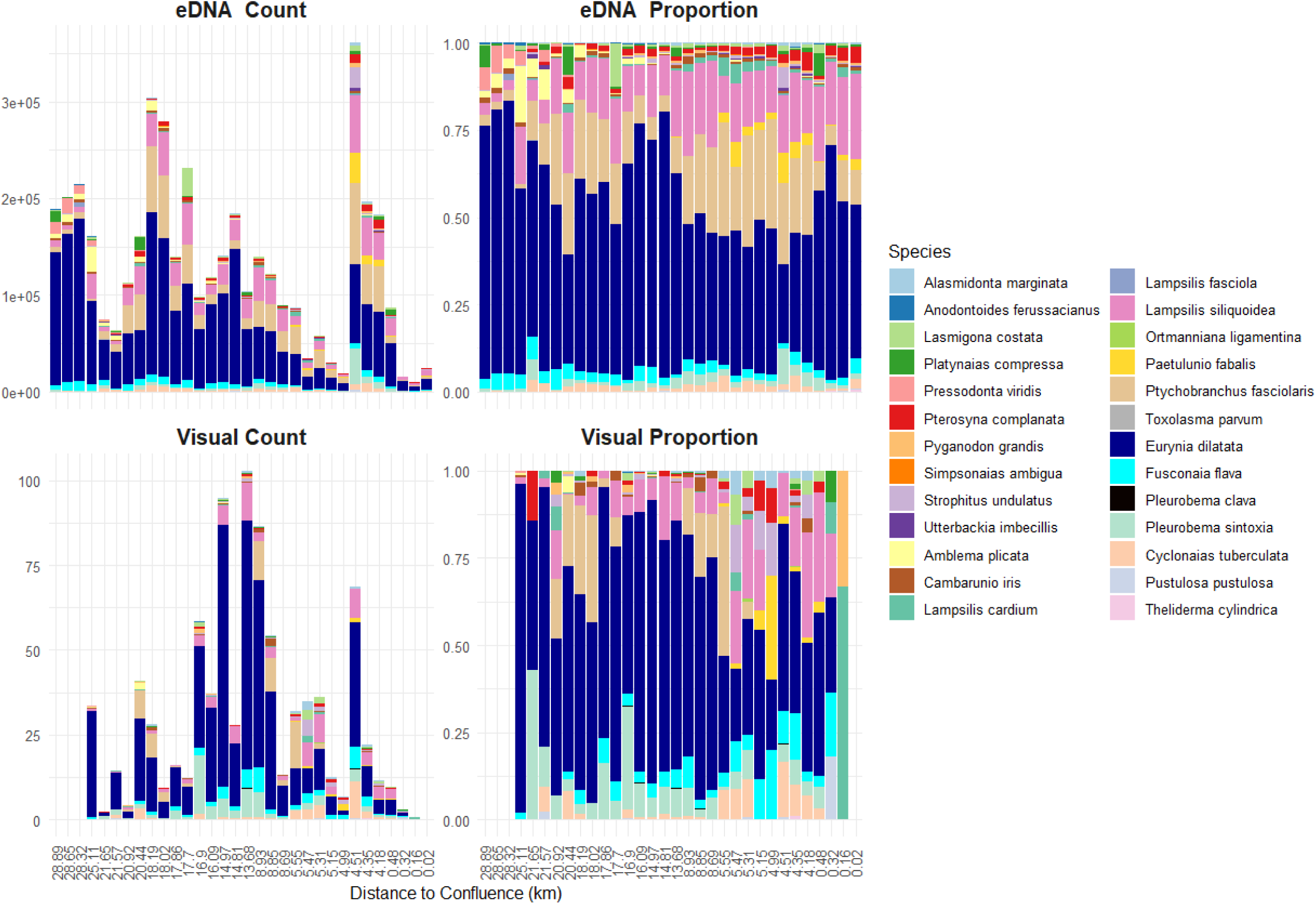
Site mussel assemblage bar plots for the number of mussels in the visual survey or the number of eDNA sequences in the eDNA survey.

**Supplementary Figure 5.**
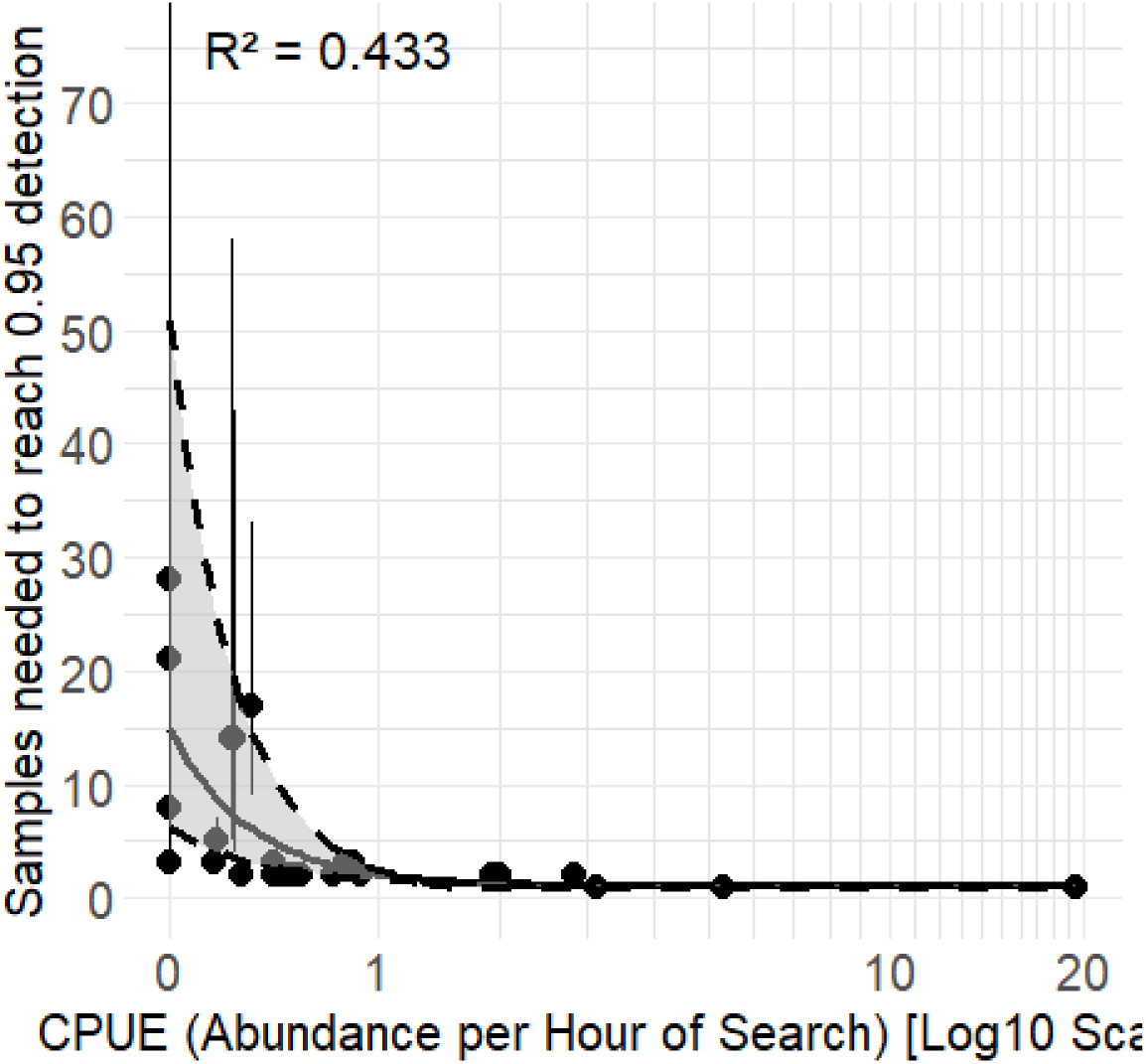
The estimated number of samples required to exceed a 0.95 probability of detection compared to the observed CPUE for each species.

**Supplementary Figure 6.**
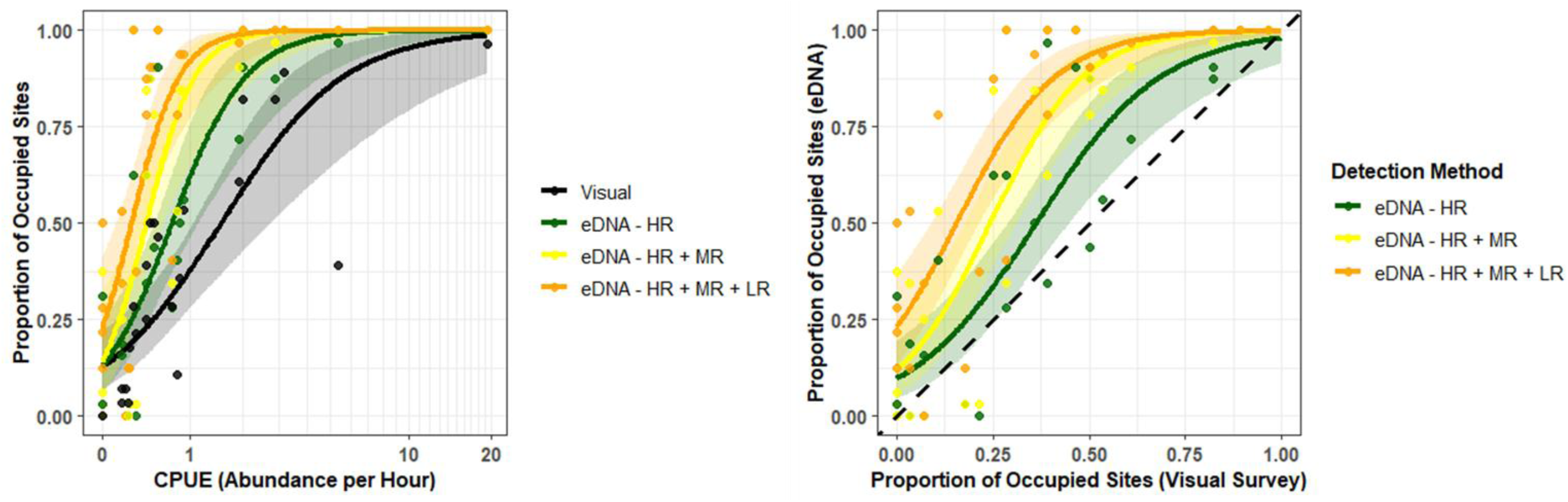
(A) Logistic regression of proportion of occupied sites from visual surveys or eDNA surveys compared t o the mussel abundance from visual surveys for all detected mussel species. (B) Logistic regression of proportion of occupied sites from eDNA surveys compared to proportion of occupied sites from visual surveys for all detected mussel species. Dashed line represents a 1:1 relationship between eDNA and visual surveys. Color curves represent eDNA detections across the repeatability scale.

**Supplementary Figure 7.**
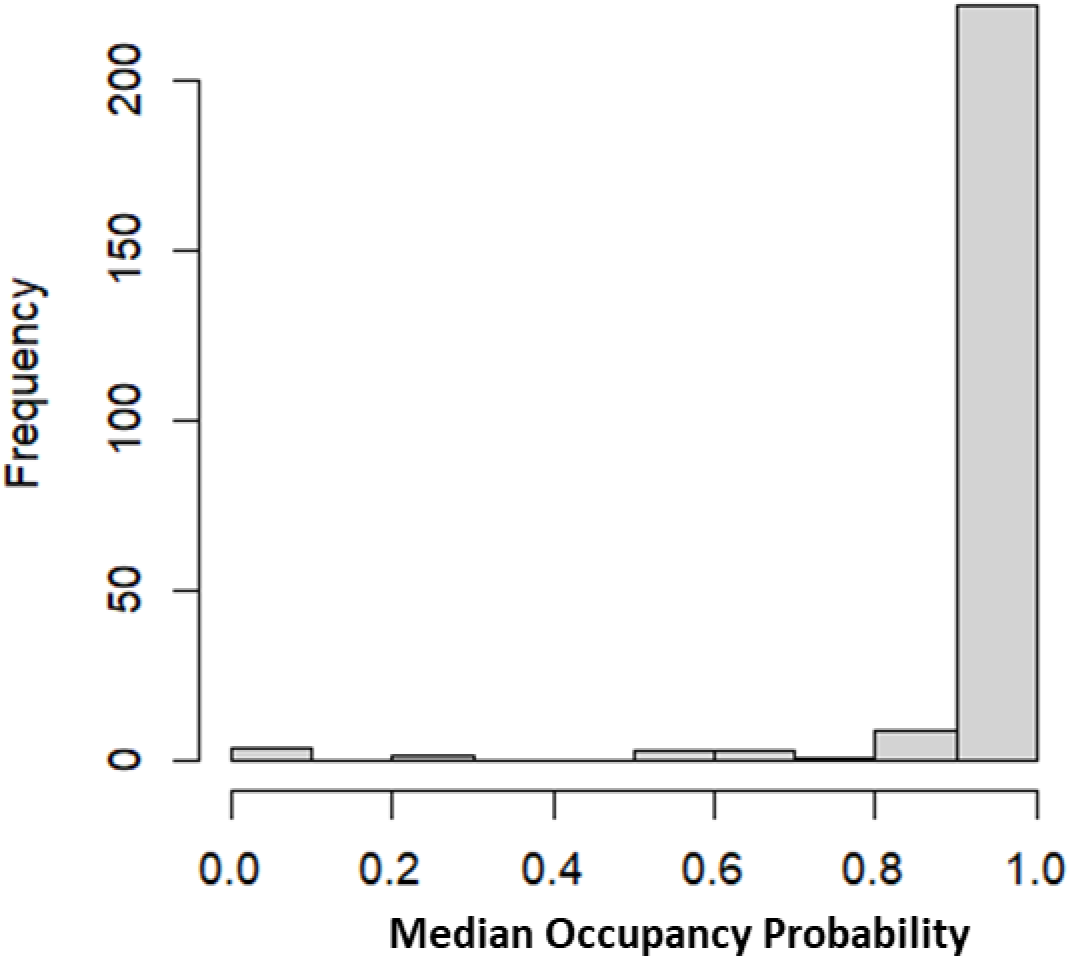
Histogram of the frequency of estimated species occupancy from eDNA data associated with visual species observations. Refer to Figure 6.

**Supplementary Figure 8.**
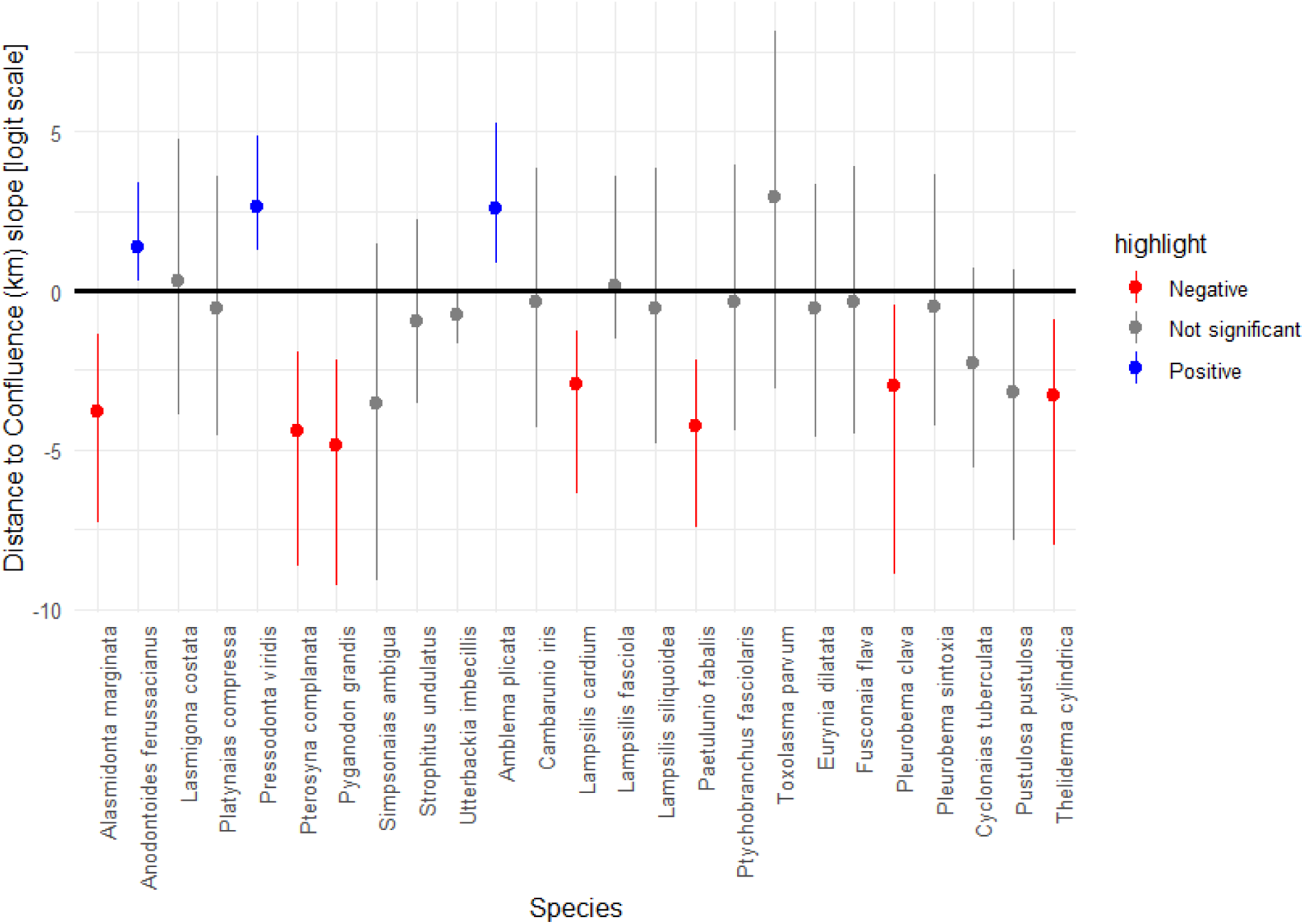
Posterior model estimates for species occupancy related to distance to the river confluence.

**Supplementary Figure 9.**
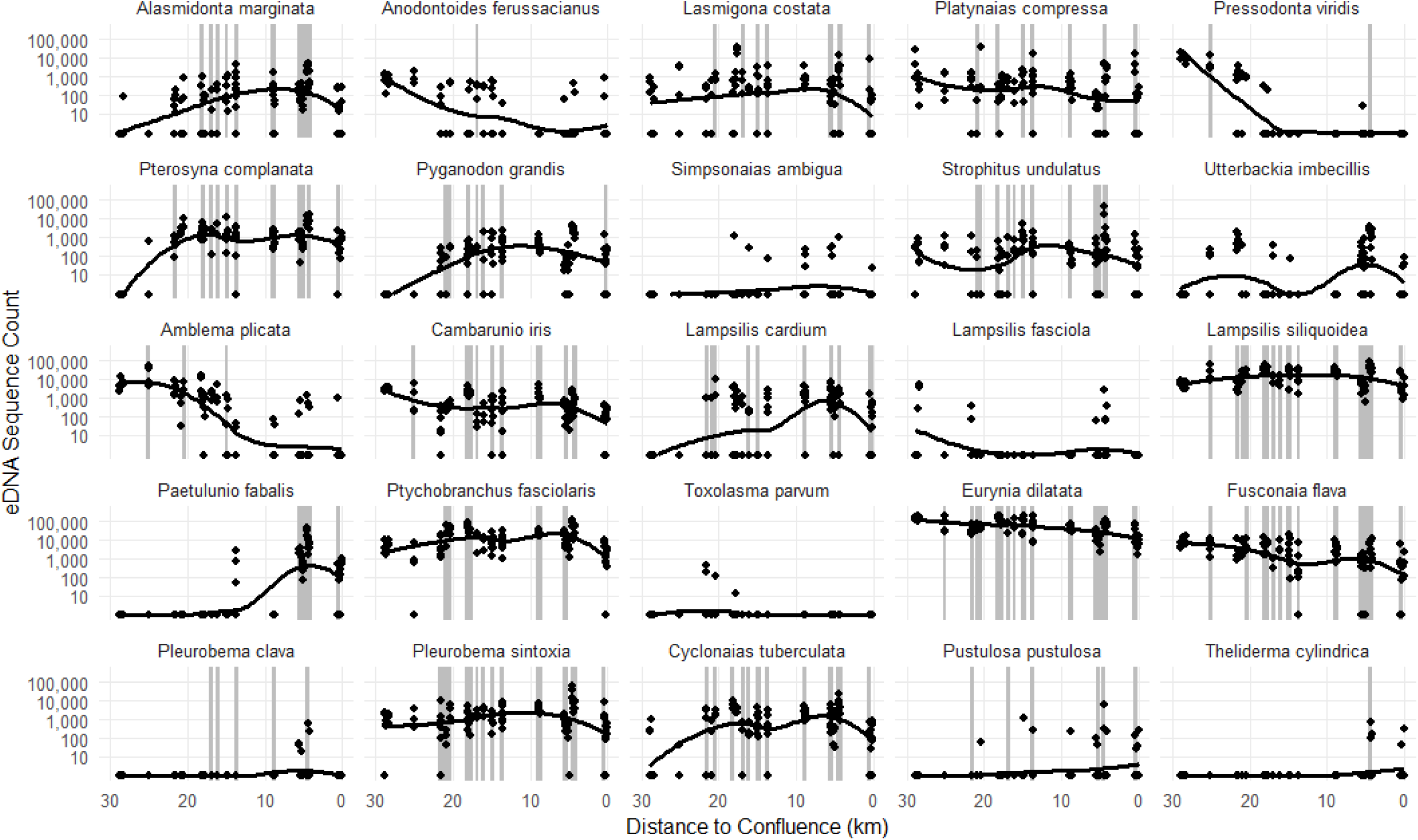
The observed eDNA sequence counts for each species across Fish Creek. Grey shading represents sites wi th visual observations for each species.

**Supplementary Table 1.**
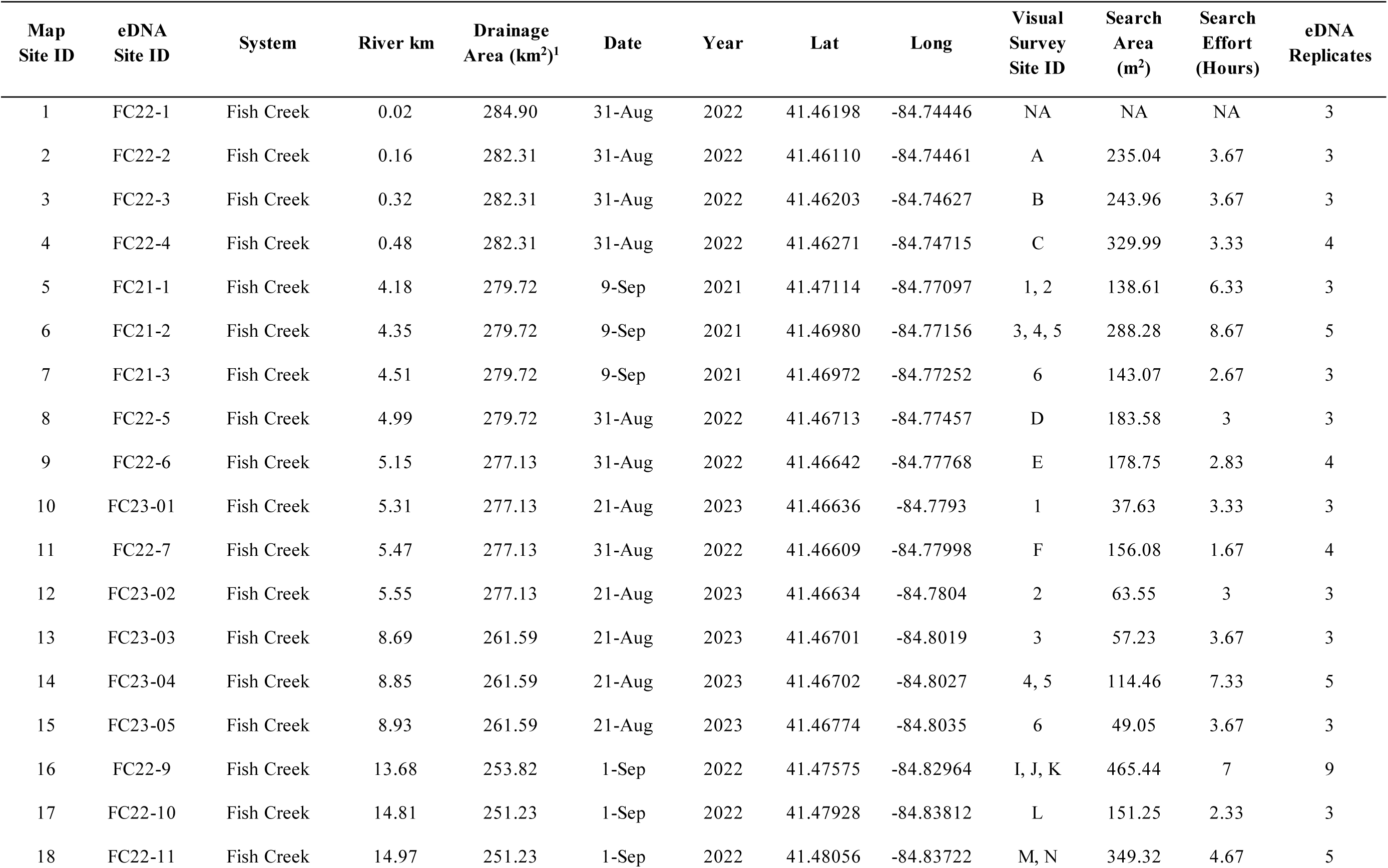
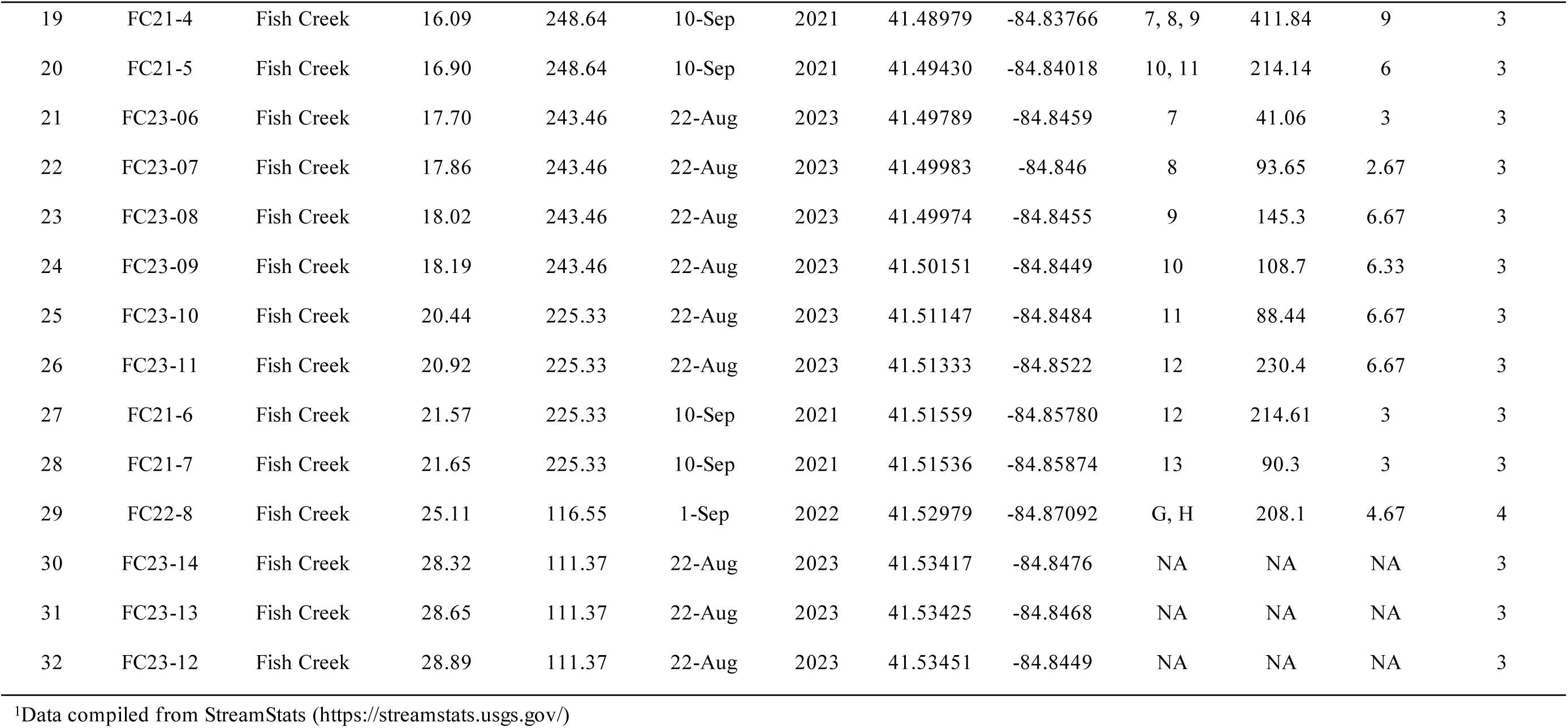
Survey site information for eDNA sampling conducted in Fish Creek and Walhonding River.

## References

Beng, K. C., & Corlett, R. T. (2020). Applications of environmental DNA (eDNA) in ecology and conservation: opportunities, challenges and prospects. Biodiversity and Conservation, 29, 2089–2121.

Bohmann, K., Elbrecht, V., Carøe, C., Bista, I., Leese, F., Bunce, M.,… & Creer, S. (2022). Strategies for sample labelling and library preparation in DNA metabarcoding studies. Molecular ecology resources, 22(4), 1231–1246. 10.1111/1755-0998.13512

Bolyen, E., Rideout, J. R., Dillon, M. R., Bokulich, N. A., Abnet, C. C., Al-Ghalith, G. A.,… & Caporaso, J. G. (2019). Reproducible, interactive, scalable and extensible microbiome data science using QIIME 2. Nature biotechnology, 37(8), 852–857.

Bylemans, J., Gleeson, D. M., Duncan, R. P., Hardy, C. M., & Furlan, E. M. (2019). A performance evaluation of targeted eDNA and eDNA metabarcoding analyses for freshwater fishes. Environmental DNA, 1(4), 402–414.

Callahan, B. J., McMurdie, P. J., Rosen, M. J., Han, A. W., Johnson, A. J. A., & Holmes, S. P. (2016). DADA2: High-resolution sample inference from Illumina amplicon data. Nature methods, 13(7), 581–583.

Camacho, C., Coulouris, G., Avagyan, V., Ma, N., Papadopoulos, J., Bealer, K., & Madden, T. L. (2009). BLAST+: architecture and applications. BMC bioinformatics, 10, 1–9.

Coghlan, S. A., Currier, C. A., Freeland, J., Morris, T. J., & Wilson, C. C. (2021). Community eDNA metabarcoding as a detection tool for documenting freshwater mussel (Unionidae) species assemblages. Environmental DNA, 3(6), 1172–1191.

Deiner, K., & Altermatt, F. (2014). Transport distance of invertebrate environmental DNA in a natural river. PloS one, 9(2), e88786.

Deiner, K., Bik, H. M., Mächler, E., Seymour, M., Lacoursière-Roussel, A., Altermatt, F.,… & Bernatchez, L. (2017). Environmental DNA metabarcoding: Transforming how we survey animal and plant communities. Molecular ecology, 26(21), 5872–5895.

Deiner, K., Yamanaka, H., & Bernatchez, L. (2021). The future of biodiversity monitoring and conservation utilizing environmental DNA. Environmental DNA, 3(1), 3–7.

Dokai, W. K., Barry, P. D., Zanatta, D. T., Gruenthal, K. M., McPhee, M. V., McIntyre, P. B., & Larson, W. A. (2023). Two for the price of one: eDNA metabarcoding reveals temporal and spatial variability of mussel and fish co-distributions in Michigan riverine systems. Environmental DNA, 5(3), 424–437.

Doser, J. W., Finley, A. O., Kéry, M., & Zipkin, E. F. (2022). spOccupancy: An R package for single-species, multi-species, and integrated spatial occupancy models. Methods in Ecology and Evolution, 13(8), 1670–1678.

Douglass, S. A., Palmer, S., McCallum, A. R., Reves, O. P., Robinson, H. A., Rutledge, A. J.,… & Davis, M. A. (2025). Environmental DNA reveals the Salamander Mussel *Simpsonaias ambigua* alive in Illinois, USA, after a century in obscurity. Ecology, 106(7), e70145. 10.1002/ecy.70145

Esling, P., Lejzerowicz, F., & Pawlowski, J. (2015). Accurate multiplexing and filtering for high-throughput amplicon-sequencing. Nucleic acids research, 43(5), 2513–2524. 10.1093/nar/gkv107

Ficetola, G. F., Pansu, J., Bonin, A., Coissac, E., Giguet-Covex, C., De Barba, M.,… & Taberlet, P. (2015). Replication levels, false presences and the estimation of the presence/absence from eDNA metabarcoding data. Molecular ecology resources, 15(3), 543–556.

FMCS Freshwater Mollusk Conservation Society. 2025. The 2025 checklist of freshwater bivalves (Mollusca: Bivalvia: Unionida) of the United States and Canada. Considered and approved by the Bivalve Names Subcommittee August 2025.

Gutiérrez, J. L., Jones, C. G., Strayer, D. L., & Iribarne, O. O. (2003). Mollusks as ecosystem engineers: the role of shell production in aquatic habitats. Oikos, 101(1), 79–90.

Haag, W. R. (2012). North American freshwater mussels: natural history, ecology, and conservation. Cambridge University Press.

Haag, W. R., & Williams, J. D. (2014). Biodiversity on the brink: an assessment of conservation strategies for North American freshwater mussels. Hydrobiologia, 735, 45–60.

Jerde, C. L. (2021). Can we manage fisheries with the inherent uncertainty from eDNA?. Journal of fish biology, 98(2), 341–353.

MacKenzie, D. I., Nichols, J. D., Lachman, G. B., Droege, S., Andrew Royle, J., & Langtimm, C. A. (2002). Estimating site occupancy rates when detection probabilities are less than one. Ecology, 83(8), 2248–2255.

Marshall, N. T., Symonds, D. E., Dean, C. A., Schumer, G., & Fleece, W. C. (2022). Evaluating environmental DNA metabarcoding as a survey tool for unionid mussel assessments. Freshwater Biology, 67(9), 1483–1507.

Marshall, N. T., & Fleece, W. C. (2025). Assessment of Environmental DNA Survey Design for the Detection of Freshwater Unionid Mussels. Environmental DNA, 7(4), e70152.

Marshall, N. T., Symonds, D. E., & Fleece, W. C. (2025). Environmental DNA detection of the male mitochondrial genome of freshwater mussels (Unionidae). Genome, 68, 1–17.

Marshall, N. T., Berg, N., Dean. C., & Fleece, W. C. (2026a). Spawning trends of freshwater mussels detected with environmental DNA. Nature Scientific Reports (In Review).

Marshall, N. T., Berg, N., Mullins, R., Stahlman, C., Dean, C. A., Sierra, M., & Fleece, W. C. (2026b). Assessment of Environmental DNA Surveys for the Cryptic Salamander Mussel (*Simpsonaias ambigua*). Biological Conservation (In Review).

Marshall, N. T., Johnson, N., Dean, C. A., & Fleece, W. C. (2026c). Use of eDNA for assessing freshwater mussel assemblage during environmental consultation. Aquatic Conservation (In Review).

Marshall, N. T., Dean. C., & Fleece, W. C. (2026d). Seasonal Environmental DNA Detection of Freshwater Mussels. Environmental DNA (In Review).

Martin, M. (2011). Cutadapt removes adapter sequences from high-throughput sequencing reads. EMBnet. journal, 17(1), 10–12.

McClenaghan, B., Fahner, N., Cote, D., Chawarski, J., McCarthy, A., Rajabi, H.,… & Hajibabaei, M. (2020). Harnessing the power of eDNA metabarcoding for the detection of deep-sea fishes. PloS one, 15(11), e0236540.

Porto-Hannes, I., McNichols-O’Rourke, K., Goguen, M., Fang, M., & Morris, T. J. (2021). Sampling protocol for the freshwater mussel *Simpsonaias ambigua* (Salamander Mussel) in Canada. Fisheries and Oceans Canada. Canadian Technical Report of Fisheries and Aquatic Sciences 3411.

Prié, V., Valentini, A., Lopes-Lima, M., Froufe, E., Rocle, M., Poulet, N.,… & Dejean, T. (2021). Environmental DNA metabarcoding for freshwater bivalves biodiversity assessment: methods and results for the Western Palearctic (European sub-region). Hydrobiologia, 848, 2931–2950.

Prié, V., Clément, L., Gaboriaud, C., Gailledrat, M., Naudon, D., Bourri, R., Lopes-Lima, M., & Valentini, A. (2025). From the Balkans to France: the threatened range-restricted freshwater mussel Unio carneus revealed as an introduced species by environmental DNA.

Ricciardi, A., Neves, R. J., & Rasmussen, J. B. (1998). Impending extinctions of North American freshwater mussels (Unionoida) following the zebra mussel (*Dreissena polymorpha*) invasion. Journal of animal ecology, 67(4), 613–619.

Richardson, R. T. (2022). Controlling critical mistag-associated false discoveries in metagenetic data. Methods in Ecology and Evolution, 13(5), 938–944. 10.1111/2041-210X.13838

Ruppert, K. M., Kline, R. J., & Rahman, M. S. (2019). Past, present, and future perspectives of environmental DNA (eDNA) metabarcoding: A systematic review in methods, monitoring, and applications of global eDNA. Global Ecology and Conservation, 17, e00547.

Sanchez, B., & Schwalb, A. N. (2021). Detectability affects the performance of survey methods: a comparison of sampling methods of freshwater mussels in Central Texas. Hydrobiologia, 848(12), 2919–2929.

Schnell, I. B., Bohmann, K., & Gilbert, M. T. P. (2015). Tag jumps illuminated–reducing sequence-to-sample misidentifications in metabarcoding studies. Molecular ecology resources, 15(6), 1289–1303. 10.1111/1755-0998.12402

Shogren, A. J., Tank, J. L., Egan, S. P., Bolster, D., & Riis, T. (2019). Riverine distribution of mussel environmental DNA reflects a balance among density, transport, and removal processes. Freshwater Biology, 64(8), 1467–1479.

Smith, D. R. (2006). Survey design for detecting rare freshwater mussels. Journal of the North American Benthological Society, 25(3), 701–711.

Sternhagen, E. C., Davis, M. A., Larson, E. R., Pearce, S. E., Ecrement, S. M., Katz, A. D., & Sperry, J. H. (2024). Comparing cost, effort, and performance of environmental DNA sampling and trapping for detecting an elusive freshwater turtle. Environmental DNA, 6(2), e525.

Stoeckle, B. C., Beggel, S., Kuehn, R., & Geist, J. (2021). Influence of stream characteristics and population size on downstream transport of freshwater mollusk environmental DNA. Freshwater Science, 40(1), 191–201.

USFWS United States Fish and Wildlife Service. (2020). Purple Cat’s Paw Pearlymussel (*Epioblasma obliquata obliquata*) 5-Year Review: Summary and Evaluation. Ecological Services Field Office Columbus, Ohio. https://ecos.fws.gov/docs/five_year_review/doc6391.pdf

USFWS United Fish and Wildlife Service (2023). Species Status Assessment Report for the Salamander Mussel (Simpsonaias ambigua). https://ecos.fws.gov/ServCat/DownloadFile/235640

USFWS (United States Fish and Wildlife Service). (2023). ECOS Environmental Conservation Online System. https://ecos.fws.gov/ecp/. Accessed in December 2025.

Van Driessche, C., Everts, T., Neyrinck, S., & Brys, R. (2023). Experimental assessment of downstream environmental DNA patterns under variable fish biomass and river discharge rates. Environmental DNA, 5(1), 102–116.

Vaughn, C. C. (2018). Ecosystem services provided by freshwater mussels. Hydrobiologia, 810, 15–27.

Watters, G. T. (1998). Freshwater mussels surveys of the Fish Creek system in Ohio and Indiana. Ohio Biological Survey Notes 1: 25–29.

Watters, G. T. (2000). Three year freshwater mussel life requirement investigation. Final report to The Fish Creek Trust Committee. Ohio Biological Survey and Ohio State University. 30 pp.

Watters, G. T., Hoggarth, M. A., & Stansberry, D. H. (2009). The freshwater mussels of Ohio. The Ohio State University Press.

Wilcox, T. M., McKelvey, K. S., Young, M. K., Sepulveda, A. J., Shepard, B. B., Jane, S. F.,… & Schwartz, M. K. (2016). Understanding environmental DNA detection probabilities: A case study using a stream-dwelling char *Salvelinus fontinalis*. Biological Conservation, 194, 209–216.

Williams, J. D., Warren Jr, M. L., Cummings, K. S., Harris, J. L., & Neves, R. J. (1993). Conservation status of freshwater mussels of the United States and Canada. Fisheries, 18(9), 6–22.

Williams, J. D., Bogan, A. E., Butler, R. S., Cummings, K. S., Garner, J. T., Harris, J. L.,… & Watters, G. T. (2017). A revised list of the freshwater mussels (Mollusca: Bivalvia: Unionida) of the United States and Canada. Freshwater Mollusk Biology and Conservation, 20(2), 33–58.

Wisniewski, J. M., Rankin, N. M., Weiler, D. A., Strickland, B. A., & Chandler, H. C. (2013). Occupancy and detection of benthic macroinvertebrates: a case study of unionids in the lower Flint River, Georgia, USA. Freshwater Science, 32(4), 1122–1135.

Yates, M. C., Glaser, D. M., Post, J. R., Cristescu, M. E., Fraser, D. J., & Derry, A. M. (2021). The relationship between eDNA particle concentration and organism abundance in nature is strengthened by allometric scaling. Molecular Ecology, 30(13), 3068–3082.

